# Eco-evolutionary dynamics of anthelmintic resistance in soil-transmitted helminths

**DOI:** 10.1101/2024.05.03.591449

**Authors:** Swati Patel, Kelsey Lyberger, Carolin Vegvari, Hayriye Gulbudak

## Abstract

Anthelmintic resistance (AR) of helminth parasites against the most widely available drugs is an ongoing concern for both human and livestock-infecting species. Indeed, there has been substantial evidence of AR in livestock but less in humans, which may be due to a variety of reasons. In this paper, we develop an eco-evolutionary model that couples the life cycle of these parasites with their underlying evolution in a single biallelic genetic locus that confers resistance to treatment drugs. We determine the critical treatment frequency needed to effectively eliminate the population, for a fixed drug efficacy (without evolution) and use this to classify three qualitative distinct behaviors of the eco-evolutionary model. Then, we describe how aspects of the life cycle influence which qualitative outcome is achieved and the spread of the resistance allele, comparing across parameterized models of human- and livestock-infecting species. For all but one species, we find that lower fecundity rates and lower contact rates speed the spread of resistance, while lower larval death slows it down. The life cycle parameters of *Ancylostoma duodenale* and *Ostertagia circumcincta* are associated with the fastest and slowest spread of resistance, respectively. We discuss the mechanistic reason for these results.

## 1 Introduction

Soil-transmitted helminths are parasitic worms and amongst the most prevalent of the neglected tropical diseases. In early 2000s, the World Health Organization began implementing mass drug administration (MDA) to combat this pervasive parasite across sub-Saharan Africa, Asia and South America, in order to meet their targets of elimination. Still, soil-transmitted helminths (STH) infect more than 1.5 billion people worldwide [1]. In many places, the current treatment strategies targets a coverage of 75% of school-age children and pre-school-age children once to twice a year, depending on endemicity levels [2]. However, there are considerations of extending treatment to adults to achieve elimination [3]. This strategy may pose an increased risk for the development of anthelmintic resistance [4]. Indeed, there has been substantial evidence of anthelmintic resistance in related parasitic helminth species that infect livestock host populations, such as sheep, goats, horses and cattle [5], and mixed evidence in human host populations [6]. It is not well understood why drug resistance evolves in some populations and not others. In this study, we develop and use a simple analytically tractable mathematical model to investigate what are the factors that influence the evolution of drug resistance in soil-transmitted helminths.

Drug resistance of related helminth parasites in livestock populations has become a significant problem, with resistance to several classes of drugs arising as early as the 1980’s [7]–[9]). Specifically, resistance became prevalent in sheep and goats, and then increased in cattle [5], [10]–[12]. Several factors likely contributed to the rapid rise of resistance in livestock including high frequency of treatment, high-coverage mass treatment, and underdosing [13].

Due to this rise of anthelmintic resistance in livestock populations, much effort has been placed on deciphering the genetic basis of resistance to the different classes of drugs (for a review see [14]). Arguably, benzimidazole resistance is best understood, in which several mutations (or single-nucleotide polymorphisms) in the *β*-tubulin genes have been identified as leading to resistance. Most of this evidence has been generated from genetic association studies, first in the model organism *C. elegans* [15] and then in the parasitic species, *H. contortus* [16] and *T. colubriformis* [17]. Still, we do not have a complete understanding of the genetics behind resistance for livestock species [14].

In humans, there is mixed evidence of anthelmintic resistance in soil-transmitted helminths, which are predominantly treated with benzimidazole drugs. Typically, empirical studies for anthelmintic resistance aim to detect it through the egg reduction ratio tests [18] or genetic screening of the SNPs that have been associated with resistance in livestock. Decreases in egg reduction ratios are thought to be indicative of reduced drug efficacy, but this measure can be biased [19]. Studies that have looked for resistance-associated SNPs have either not found them [20]–[22] or have found them at low frequencies in human helminths [23]–[26]. Additionally, the effects of these specific mutations on parasite fitness has not been thoroughly investigated.

Anthelmintic resistance models have been developed for both human and veterinary helminth parasites. Previous modeling work for livestock populations has put forth solutions for reducing anthelmintic resistance, including rotation between drugs, appropriate dosing, maintaining refugia, and rotational grazing [27]–[29]. However, such strategies may not always be feasible for human populations because of fundamental ethical and logistical constraints. Additionally, the possibility of development of resistant worm populations in humans is particularly worrisome as they are treated almost exclusively with a single drug class, benzimidazoles. In order to apply what was learned from livestock to prevent the development of resistance when treating human STH [4], [30], we must consider differences between the two systems including the underlying biology of hosts and parasites.

Modeling studies for human-infecting parasites have focused on the question whether certain characteristics of helminth population biology, such as overdispersion and positive and negative density dependence, affect the rate of evolution of resistance [31]–[33]. Parasite overdispersion is the observation that a few hosts have very high worm burdens while most hosts have few or no worms. The mating probability between male and female parasites is subject to positive density dependence because male and female parasites need to be together in the same host to reproduce. Negative density dependence can affect multiple processes in the parasite life cycle, e.g. fecundity, mortality and parasite establishment within the host. The consensus now is that overdispersion on its own does not affect the rate of spread of resistance [33], [34]. In contrast, density-dependent fecundity can speed up the rate of spread of resistance. The same has been found for density-dependent mortality and parasite establishment within hosts but to a lesser degree [33].

Here, we develop and analyze a pulsed model for drug resistance evolution in soil-transmitted helminths. We deliberately choose a model structure in which we ignore overdispersion and density-dependent processes to focus on the impact of the values of other fundamental parameters on the spread of disease and resistance. We parameterize our model with data on three human and three livestock parasitic helminth species and then use this first to ask how much drug treatment is needed for human and veterinary parasite species to reach elimination and how this depends on the drug efficacy (which may be decreasing due to resistance). Then, we compare across species to examine how the timing and coverage of drug administration, the parasite biology and the host biology impact the population dynamics of the disease. Finally, we study how these three aspects impact the patterns of resistance evolution.

## 2 Model

Soil-transmitted helminths have a multi-stage life-cycle, which is partly spent within a human host (juvenile and adult worms) and partly in an external soil environment (eggs, larvae). Building off of an established disease transmission model that incorporates the population biology of these worms, we let *W* (*t*) be the total number of worms living in a human host population of size *N* (assumed to be constant), and *L*(*t*) be the number of active larvae in the environment, at time *t* (Figure 1) [35]. Disease transmission depends on the host contact with soil and uptake of the larvae from the environment. We assume each individual host has a contact and uptake rate of *β*. We assume that the transition from larvae to adults occurs immediately upon uptake within the host (ignoring time delays for development). Additionally, adult worms have a natural intrinsic per-capita death rate, *µ_w_*, eggs and larvae have a natural intrinsic death rate, *µ_l_*, and human and livestock hosts die at a rate of *µ_h_*. Host death implies that all worms within the host also die. Adult worms produce and secrete eggs at a rate *λ*, into the soil environment.

**Figure 1:**
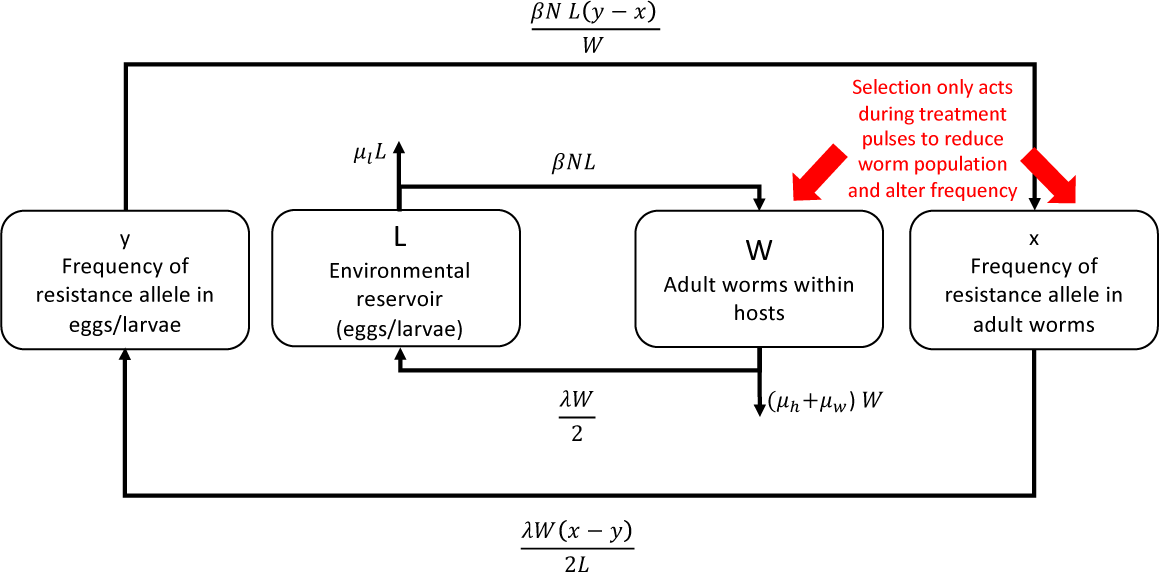
Schematic representation of our model. Larvae (*L*) in the soil get taken up by hosts and then turn into adult worms (*W*). Adult worms release eggs that turn into larvae in the soil environment. Populations are genetically structured and *x* and *y* are the frequencies of the resistance allele in the hosts and environment, respectively. Pulsed treatments (red arrows) affect the worm population and frequency of the resistance allele in the hosts.

**Figure 2:**
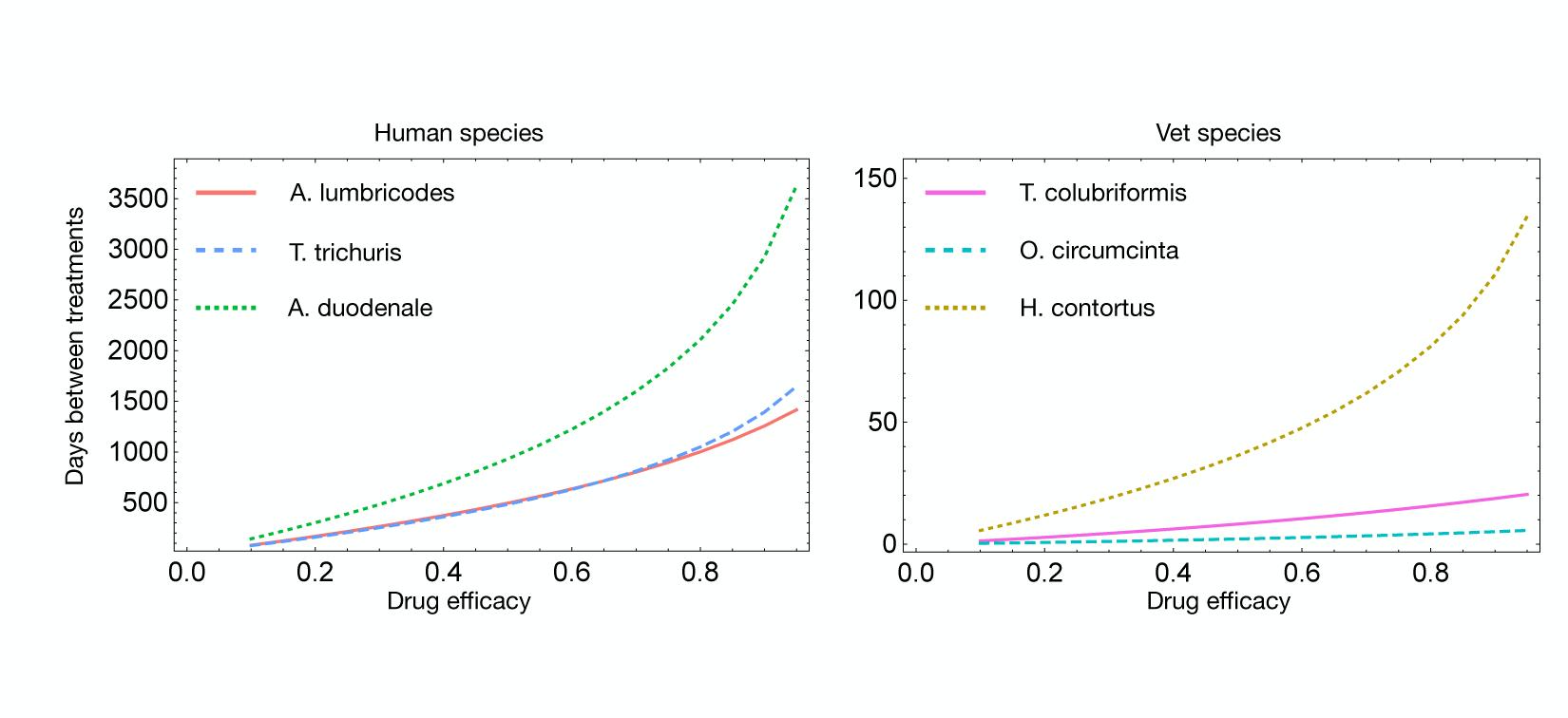
Critical time between pulses as it depends on drug efficacy in three human (A) and livestock (B) infecting parasites. Parameters are as given in Table 1 and 2 and coverage was assumed to be 1. Treatment applied less frequently than the critical time will allow parasite populations to grow, while treatment applied more frequently will lead to elimination.

Additionally, we assume the parasite populations are subject to a pulse disturbance, periodically every *τ* time units, from the mass drug administration (MDA). At the time of (MDA), some proportion of the adult worms within the host population dies, depending on the coverage (*c*) and drug efficacy (the average proportion of the worms that die within a treated host). Furthermore, we assume that there is a single locus in the worm genome that determines how it responds to the drug, i.e., the drug efficacy. We assume the locus has two possible alleles, *A* and *B*, and that *A* is a “resistance” allele, meaning it makes the worm less vulnerable to the effects of the drug. Soil-transmitted helminths are diploid organisms and we let *µ_AA_*, *µ_AB_*, *µ_BB_* represent the drug susceptibility of worms with genotype *AA, AB, BB*, respectively. We assume *µ_AA_ ≤ µ_AB_ ≤ µ_BB_*. Worms are identical in all other ways except for their genotype-dependent drug susceptibility. We let *x*(*t*) and *y*(*t*) be the frequency of the resistance allele *A*, in the hosts and soil environment, respectively.

We assume Hardy-Weinberg equilibrium so the frequency of each genotype can be determined from the frequency of alleles. Then, we let *Ū* be the average susceptibility of worms to the drug, which depends on the frequency of each genotype and hence, *x*(*t*). Dropping the dependence on *t*:

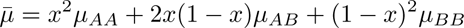

Additionally, we let 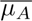 be the marginal average death rate associated with the resistance allele. This can be interpreted as the proportion of worms with at least one resistance allele that die in a treated host. This depends on *x*(*t*):

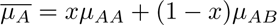

More details on these derivations can be found in the Supplement.

Altogether, this leads us to a set of impulse ordinary differential equations, describing the change in the number of adult worms within the hosts, *W* (*t*), and larvae *L*(*t*), as well as the frequency of the *A* allele (resistance) amongst the adult worm population in the host *x*(*t*) and the larvae population in the environment *y*(*t*). The dynamics are governed by:

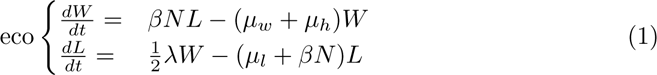

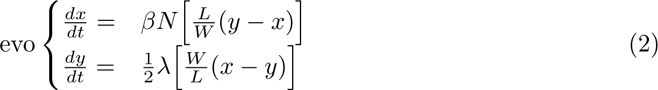

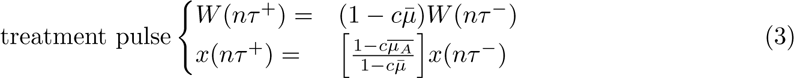

The last two equations capture the instantaneous effect of the pulsed disturbance on the adult worm population. We refer to the dynamics of *W* and *L* as the “ecological dynamics”, since they capture the abundance of worms and larvae. We refer to the dynamics of *x* and *y* as the “evolutionary dynamics” since they capture the changes in frequency of genotypes. Our model is a linear model with impulses from the treatment. Hence, it is a simple model, which makes it tractable and provides insights into the spread of disease and anthelmintic resistance.

## 3 Methods

We use this baseline model to understand how treatment regime (frequency and coverage), host properties (contact rate, *β* and host density, *N*) and parasite biology (larval death rate, *µ_l_*, adult worm death rate, *µ_w_* and fecundity, *λ*) impact the spread of disease and the evolution of resistance. We use a combination of analytical and numerical techniques to do this. Our results are divided into three parts. In the first part, we determine a critical treatment frequency, for fixed drug efficacy (with no evolution). Treating more frequently then the critical treatment frequency leads to elimination of the worm population. We use this critical frequency to distinguish three qualitatively distinct possible behaviors of the model, with respect to the spread of disease. In the second part, we analyze the sensitivity of these qualitative behaviors to three categories of parameters (treatment regime, host properties, and parasite biology). For this we use parameterizations of six different parasitic helminth species from the literature. In the third part, we investigate the sensitivity of the speed of evolution to these parameters. Numerical solutions of the model were obtained using deSolve in R [36] and code is available here: [37]

### 3.1 Parameterization and Sensitivity

To understand the effects of parameters on the eco-evolutionary dynamics, we choose values for each of the parameters from the literature separately for three different human-infecting parasite species and three veterinary parasite species (Table 1 and 2). Each species of helminth has different life history parameter values. For the human-infecting species life history parameters, we use two sources: [38], [39]. Other aspects, such as host density and treatment regimes were chosen based on specific populations. For the livestock-infecting species, we use published values from [28]. More details on how we translate parameters from these sources into our model is given in the supplement. For the ecological sensitivity analysis, we conduct a pairwise parameter sensitivity analysis, varying two parameters at a time, while fixing the others at species-specific parameter values as reported in the literature. The ranges of values were chosen to encompass the parameterized values of all six species. For the evolutionary sensitivity analysis, we choose three values of the parameter of interest, corresponding to an *R*_0_ (see eq 4) of 1, 5, or 10, and fix all other parameters at their species-specific parameter values. Since genotype-dependent drug efficacy values are not empirically well-established, we assume *µ_AA_* = 0.1*, µ_AB_* = 0.5, and *µ_BB_* = 0.9. We solve the model using these parameters to compare the effect of each parameter on the speed of resistance evolution.

**Table 1:** Parameter values used for of three human-infecting soil-transmitted helminth species. Details on the sources and derivations for these parameters are given in the supplement [38]–[40]. The ratio of worms to larvae is derived from these parameters from the expression 5 in the main text.

| Parameter | Description | *A. lumbricoides* | *A. duodenale* | *T. trichiura* |
| --- | --- | --- | --- | --- |
| $N [m^{-2}]$ | population density | $1.2 \times 10^{-4}$ | $1.2 \times 10^{-4}$ | $1.2 \times 10^{-4}$ |
| $\beta [m^2 day^{-1}]$ | contact rate | $1.1 \times 10^{-4}$ | $9.6 \times 10^{-4}$ | $1.7 \times 10^{-3}$ |
| $\lambda [day^{-1}]$ | egg production rate | $1 \times 10^4$ | $3 \times 10^3$ | $2 \times 10^3$ |
| $\mu_w [day^{-1}]$ | adult worm death rate | 0.0027 | 0.0013 | 0.0027 |
| $\mu_l [day^{-1}]$ | larvae death rate | 0.016 | 0.082 | 0.05 |
| $\mu_h [day^{-1}]$ | host death rate | $4.6 \times 10^{-5}$ | $4.6 \times 10^{-5}$ | $4.6 \times 10^{-5}$ |
| | Ratio of worms to larvae | $3.5 \times 10^{-6}$ | $5.5 \times 10^{-5}$ | $5.1 \times 10^{-5}$ |

**Table 2:** Parameter values used for livestock-infection helminth species. Details on the source [28] and derivations for these parameters are given in the supplement. The ratio of worms to larvae is derived from these parameters from the expression 5 in the main text.

| Parameter | Description | *T. colubriformis* | *O. circumcincta* | *H. contortus* |
| --- | --- | --- | --- | --- |
| $N \text{ [m}^{-2}\text{]}$ | host density | 0.001 | 0.002 | 0.001 |
| $\beta \text{ [m}^2\text{days}^{-1}\text{]}$ | contact rate | 3.08 | 2.62 | 2.44 |
| $\lambda \text{ [days}^{-1}\text{]}$ | egg production rate | 6.75 | 4.8 | 15.07 |
| $\mu_w \text{ [days}^{-1}\text{]}$ | adult worm death rate | 0.05 | 0.041 | 0.01 |
| $\mu_l \text{ [days}^{-1}\text{]}$ | larvae death rate | 0.078 | 0.03 | 0.63 |
| $\mu_h \text{ [days}^{-1}\text{]}$ | host death rate (sheep) | $2.7 \times 10^{-4}$ | $2.7 \times 10^{-4}$ | $2.7 \times 10^{-4}$ |
|  | Ratio of worms to larvae | 0.0035 | 0.045 | 0.086 |

## 4 Results

### 4.1 Critical Frequency and Three Qualitatively Distinct Dynamics

We begin with preliminary results of the model with no treatment. In this case the ecological (transmission) dynamics do not depend on the evolutionary variables. The basic reproduction number *R*_0_ in transmission models of helminth parasites is defined as the average number of female offspring produced by one adult female worm that themselves infect a new host and reach sexual maturity [35]. With our model the equation for *R*_0_ is:

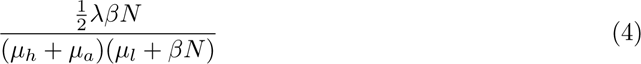

Because of the linearity of the ecological dynamics (*W* and *L*), in the absence of any treatment pulses, the worm and larvae densities either grow or decrease exponentially. If *R*_0_ *>* 1, the worm population grows. If the inequality is reversed, then the worm population decays to zero, which is consistent with an eigenvalue analysis of the linear transmission model. As expected, we observe that if the host natural death rate, adult worm death rate or the larval death rate are small, then the worm and larval populations tend to grow. Alternatively, if the fecundity is small, then the population tends to decline.

Additionally, in the long run, the worm to larvae ratio (from examining the appropriate eigen-vector) will approach

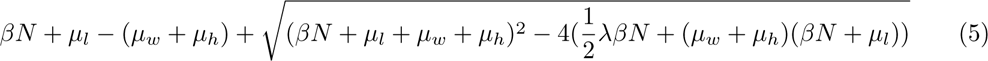

which affects the evolutionary dynamics. We compute this long term worm to larvae ratio for each species’ parameter set. Interestingly, we observe that the human-infecting species tend to have a much smaller worm to larvae ratio (most are *O*(10*^−^*^5^)) than the livestock-infecting species (most are *O*(10*^−^*^2^); see Table 1 and 2).

In the case of worm population growth, we find that, for a fixed drug efficacy (no evolution) and coverage, there is a critical frequency to treat the host population, denoted *τ ^∗^* (see supplement for mathematical derivation of an expression that gives the critical frequency and [41] for proof of existence). If treatment is administered more frequently than this critical frequency, then the worm and larvae dynamics will eventually decay to 0 as a consequence of the treatment. Conversely, if treatment is administered less frequently, then the populations will continue to grow despite repeated treatment rounds.

Using the parameter values from the literature, we numerically solve for the critical frequency of treatment needed for elimination, for any given drug efficacy and fixed coverage (code provided in the Mathematica notebook here: [37])

We find that much more frequent treatment is required to eliminate the livestock-infecting parasite species than the human-infecting species, for any given drug efficacy. For example, for a drug efficacy of 0.9, the critical time between treatments needed for human-infecting species *A. lumbricoides, A. duodenale* and *T. trichuris* is 1256, 2922, and 1393 days, respectively. On the other hand, for livestock-infecting species *T. colubriformis, O. circumcinta*, and *H. contortus* is around 18, 5, and 110 days, respectively. One key difference between livestock- and human-infecting species is that livestock-infecting species tend to have a much greater contact rate between eggs and larvae in the environment and hosts (see Table 1 and 2). Livestock species tend to live on pastures where they are also defecating.

Each treatment administration poses a selection event that instantaneously increases the frequency of the resistance allele in the host population. This decreases the average drug efficacy, which then decreases the critical time between treatments needed to drive the worm population to elimination. (Recall that each drug efficacy is associated with a critical time between treatments.) Hence, for a fixed treatment frequency, there are three qualitatively different behaviors of the ecological dynamics. First, if the time between treatments is below the critical times both before and after the evolution of resistance, then the worm population is driven to elimination even with the evolution decreasing the drug efficacy (purple curve, Figure 3). On the other hand, if the time between treatments is above the critical time even before any evolution, then the worm population grows (yellow curve, Figure 3). Finally, if the time between treatments is below the critical time for the initial drug efficacy but below for the drug efficacy after evolution, then the worm population initially decays but then begins to grow (green curve, Figure 3). This is a classical evolutionary rescue U-shaped curve [42].

**Figure 3:**
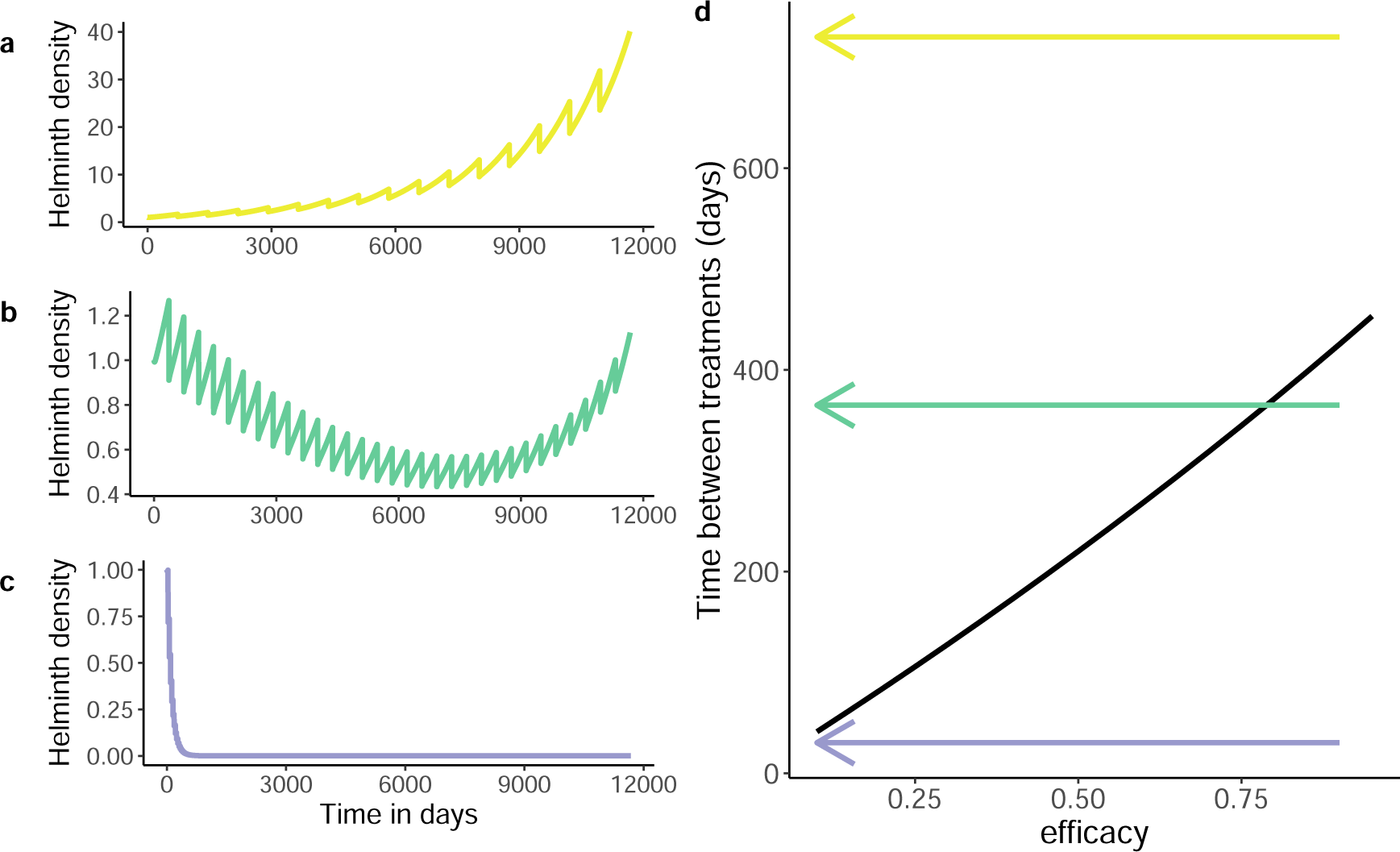
Three qualitatively distinct dynamics. Panels a-c show time series of helminth population density in (a) growth, (b) rescue, and (c) decline scenarios. *A. duodenale* parameters are used. Treatment is applied at frequencies of every 2 years, 1 year, or 1 month in a, b, and c respectively. Coverage is 0.3 to represent the goal of the WHO to reach 75 percent coverage of school-age children, which is roughly 30-40 percent of the total population in a typical endemic country. d) Critical treatment frequency threshold (expressed in time between treatments in days on the y-axis) for hookworm depending on drug efficacy. On the left of the black line the worm population grows, on the right of the black line the worm population declines. The arrows illustrate how for the same treatment frequency, reducing treatment efficacy can make the worm population shift from decline to growth (rescue scenario, green arrow) or stay within the decline (purple arrow) or growth space (yellow arrow).

Since there is no cost to resistance, eventually the frequency of the resistance allele approaches one, as selection continues through pulsed treatments. Hence, for a given set of parameters, we determine which qualitative behavior ensues, by comparing the treatment regime with the critical treatment time in the fully susceptible, denoted 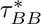 and the fully resistant populations, denoted 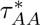. Namely, if 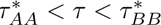, then we get evolutionary rescue. The Mathematica notebook that includes code for this categorization of the qualitative behaviors is provided here:[37].

### 4.2 Effects of Treatment Regime, Host Properties, and Parasite Biology on Qualitative Dynamics

We partition key parameters of our model into three categories, with two parameters each:

1. Treatment regime: coverage and treatment frequency
2. Host properties: *β* and *N*
3. Parasite biology: *µ_l_* and *λ*

Sensitivity patterns for *µ_w_*, the adult worm death rate, are given in the supplement. Note that *µ_h_*, the host death rate, is not considered in the analysis since it is much slower than *µ_w_* and as a result its effect is negligible.

For each category, we determine how the sensitivity of the qualitative dynamics on the two parameters over a range of parameter values, keeping other parameters fixed at their species-specific values. Additionally, we kept the genotype-dependent drug efficacies, *µ_AA_, µ_AB_, µ_BB_*, fixed across all species. We present the results for two species, *A. duodenale* as a representative of the human-infecting species, and *H. contortus*, as a representative of the livestock-infecting species. Results for other species are shown in the supplement.

#### 4.2.1 Treatment Regime

Overall, we find that increasing treatment frequency and coverage has the expected effect; more intense (higher coverage and/or higher frequency) treatment makes the population more likely to decline even with resistance evolution. For the human-infecting hookworm species, *A. duodenale*, an evolutionary rescue pattern occurs for low frequency and high population-wide coverage (e.g. 75% coverage 2x a year) or for high frequency and low coverage (e.g. 25% coverage 8x a year) treatment regimes. For high coverage and high frequency of treatment (e.g. 75% coverage 8x a year), the worm population would decline despite the evolution of resistance (Figure 4A). A similar pattern holds for the other human-infecting species, *A. lumbricoides* and *T. trichuris* (Supplement Figure S1). For the livestock-infecting species, *H. contortus*, evolutionary rescue occurs for much higher coverage and higher frequency (e.g. 75% coverage 10x a year; Figure 4B), than for hookworm. A similar pattern holds for the other livestock-infecting species, *T. colubriformis* and *O. circumcincta* (Supplement Figure S1).

**Figure 4:**
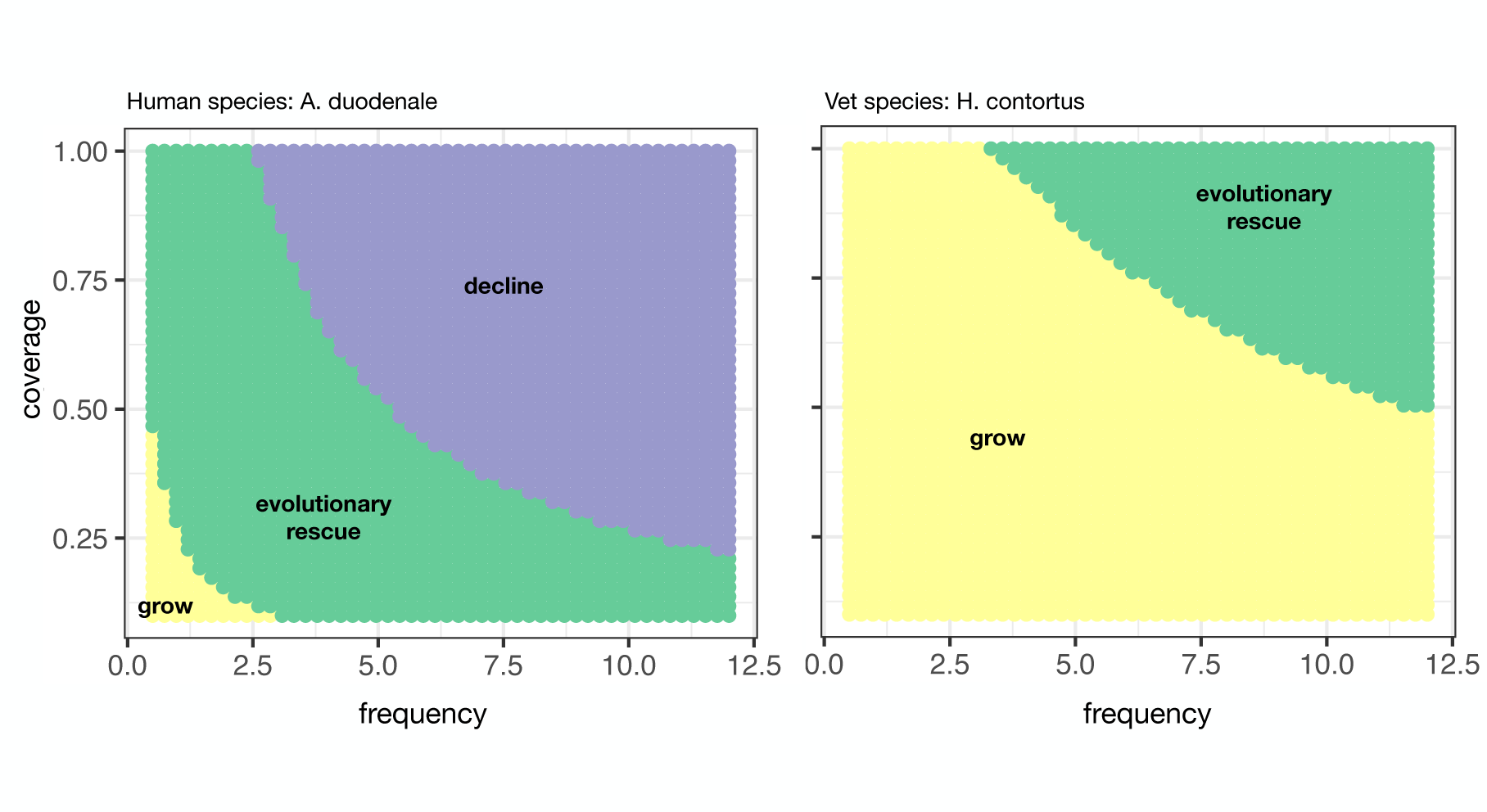
Sensitivity of qualitative dynamics on treatment parameters, frequency (number of treatments per year) and coverage (proportion of total population treated). Frequency ranged from 0.5 to 12 and coverage ranged from 0.1 to 1. Parameters for *A. duodenale* and *H. contortus* are taken from Table 1 and 2.

#### 4.2.2 Host Properties

For host properties, we find that decreasing contact rate and host density make the worm population more likely to decline to elimination despite the evolution of resistance. For *A. duodenale*, contact rates and host densities must be very small to avoid growth of the worm population (Figure 5A). On the other hand, for *H. contortus*, the contact rates and host densities can be relatively high, and the worm population still declines to elimination. For *H. contortus*, evolutionary rescue occurs for intermediate contact rates and host densities (Figure 5B). Note, for a coverage of 1 and treatment once a year, the species-specific parameters (star in Figure 5) indicate that *H. contortus* will grow, even with a fully susceptible worm population, and *A. duodenale* will undergo evolutionary rescue. Further, Figure 5 highlights the importance of the contact rate as a key differences between livestock- and human-infecting species. All else equal, the livestock-infecting parasites need a much higher contact rate with hosts to grow than human-infecting parasites.

**Figure 5:**
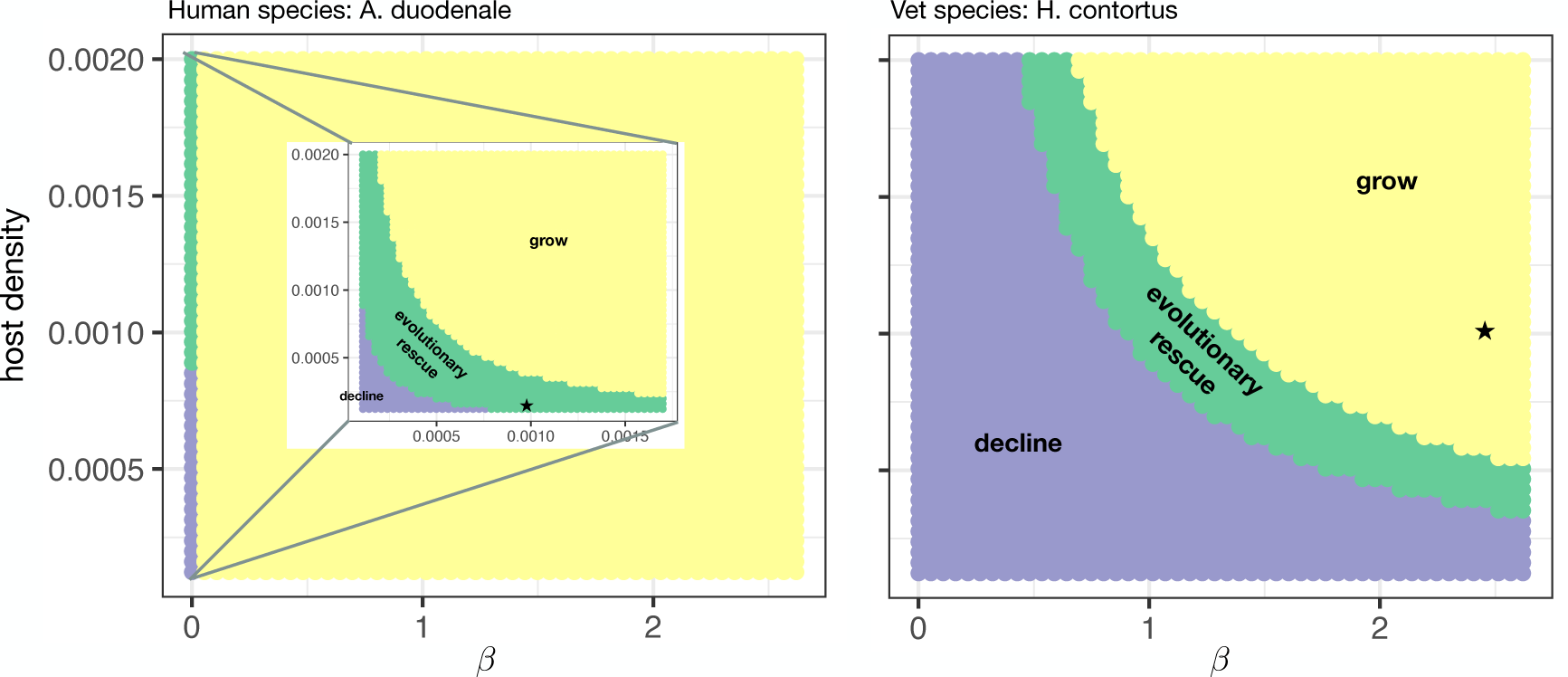
Sensitivity of qualitative dynamics on host parameters, *β* (contact rate) and host density (individuals per *m*^2^). *β* ranged from 0.00011 to 0.62 and host density from 0.00012 to 0.002. Frequency is once a year. Coverage is 1. All other parameters for *A. duodenale* and *H. contortus* are taken from Table 1 and 2. The inset is given for a smaller range of *β* from 0.00011 to 0.0017, encompassing parameter values for human helminth parasite species. Stars indicate the parameter values for *β* and host density reported for each species in the literature.

#### 4.2.3 Parasite Biology

Finally, for parameters describing parasite biology, we find that lower larval death rate and fecundity make the worm population more likely to decline, despite the evolution of resistance. For *A. duodenale*, the worm population will decline for a wide range of larval death rate and fecundity parameters and the evolutionary rescue pattern is only expected over a narrow range of high fecundity and low larval death rates. On the contrary, for *H. contortus* we find that the population will grow over a wide range of fecundity and larval death rates, even in the absence of evolution. Only over a very small range with high larval death rates and very low fecundity do we expect evolutionary rescue (Figure 6B).

**Figure 6:**
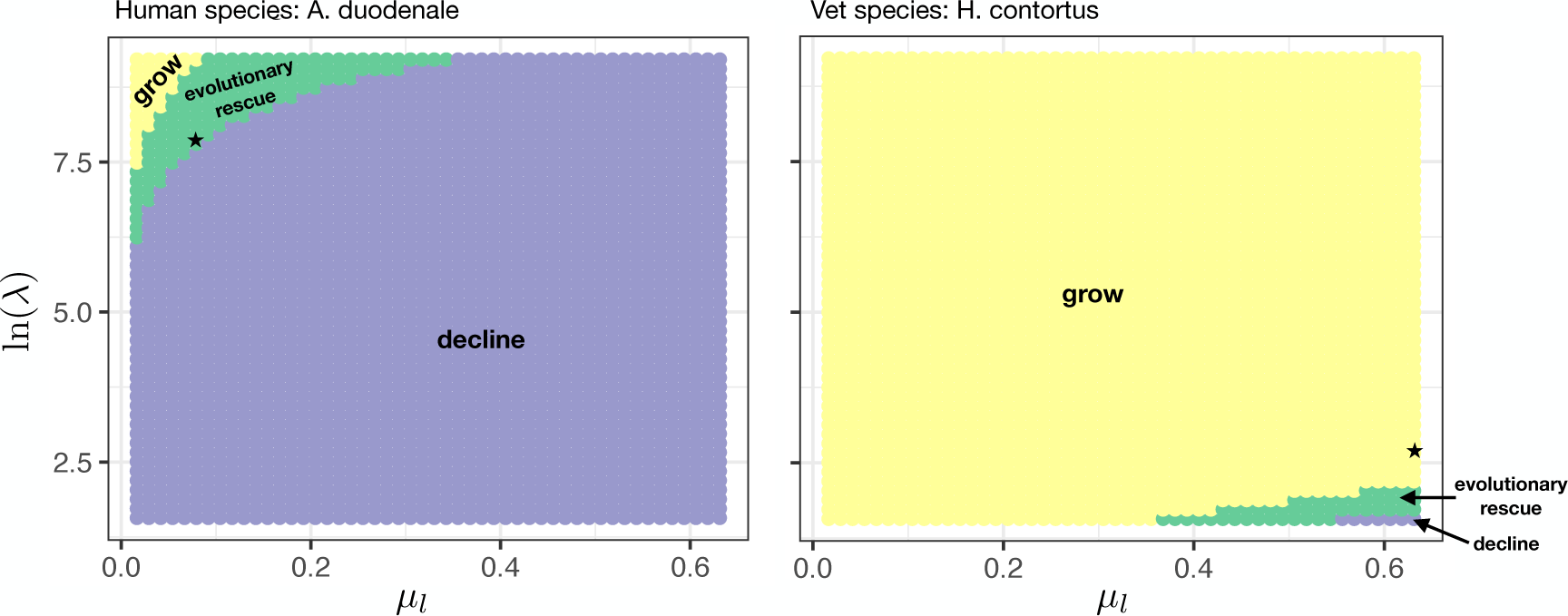
Sensitivity of qualitative dynamics on parasite parameters, *µ_l_*(larval death rate) and *λ* (egg production rate). *µ_l_* ranged from 0.016 to 0.63 and *λ* from 4.8 to 1000. Frequency is once a year. Coverage is 1. All other parameters for *A. duodenale* and *H. contortus* are taken from Table 1 and 2. Stars indicate the parameter values for *µ_l_* and *λ* for each species reported in the literature.

### 4.3 Effects of Parasite Biology, Host Properties and Treatment Regime on the Resistance Evolution

#### 4.3.1 Parasite Biology and Host Properties

The parasite biology and host properties of different parasitic helminth populations impact the patterns of drug resistance evolution, i.e., the dynamics of the frequency of the resistance allele. In particular, these parameters impact how quickly parasite populations will evolve resistance. We find that, in our model, the general pattern of drug resistance evolution occurs in two steps: (1) at the time of the pulsed treatment, there is a selection event that causes an immediate jump in the frequency of the resistant allele by disproportionately killing off the more susceptible types and (2) the jump is followed by a relatively slower decay in the frequency of the resistance alleles, due to an influx of both susceptible and resistance alleles from the soil environment, which was not directly influenced by the pulsed selection event. Together the immediate jump in frequency of resistance and the decay in the frequency of resistance alleles affect how quickly resistance evolves.

The initial jump is only influenced by the preceding resistance frequency, the coverage, and the drug efficacy (see equation in model). On the other hand, the subsequent decay depends directly on the contact rate, *β*, the host density, *N*, the fecundity, *λ*, and the relative abundance of worms to larvae (which depends also on other parameters; see equation 5). The greater the worm to larvae ratio, the slower the decay of the resistance frequency following a selection event, and hence, the faster the evolution of resistance. If the worm to larvae ratio is fixed, then greater *β* and *N* lead to faster decay (see equation 3). In contrast, greater *λ* leads to a slower decay if the worm to larvae ratio is fixed. However, *β, N,* and *λ* all indirectly affect the worm to larvae ratio. Hence, how these parameters influence the evolution, through these direct and indirect effects, is not obvious.

Comparing across species, we find that for a fixed treatment regime, human-infecting species tend to evolve resistance more quickly than livestock-infecting species, except for *A. lumbricoides* (see Figure 7). To better understand what aspects of each parasite species influence resistance evolution, we compared trajectories of the frequency of the resistant allele in the adult population, varying the value of one parameter at a time. In particular, we examined how fecundity, parasite adult and larval death rates, and contact rate of hosts with the environmental reservoir of eggs/larvae influence the two step evolutionary process. Figure 8 shows the trajectories of the frequency of the resistance allele within the host (*x*(*t*)) for *A. duodenale* and *O. circumcincta*. Trajectories of the other species show similar patterns to *A. duodenale* (Supplement Figure S2).

**Figure 7:**
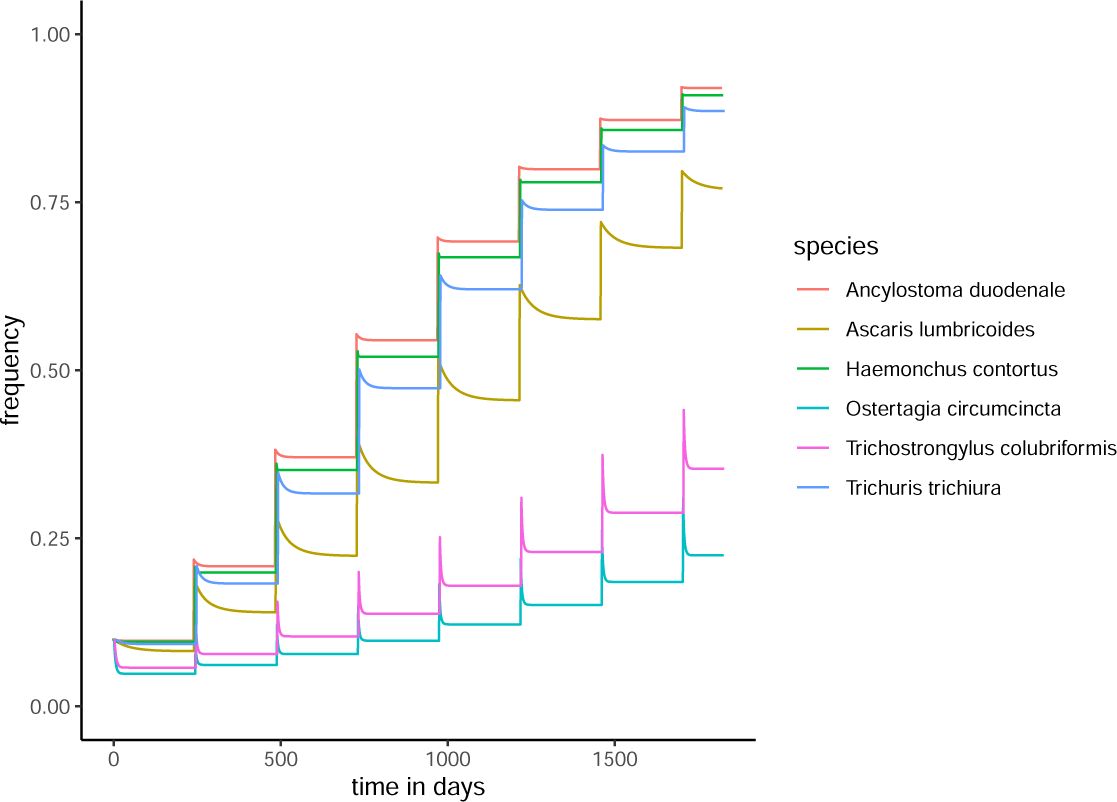
Evolutionary trajectories of *x*(*t*) for fixed treatment frequency (1*/τ* = 1.5) and coverage (*c* = 1) for six different soil-transmitted helminth parasite species. Parameter values of each species are given in Table 1 and 2. Genotype-dependent drug efficacies are fixed to be *µ_AA_*= 0.1*, µ_AB_* = 0.5*, µ_BB_* = 0.9.

**Figure 8:**
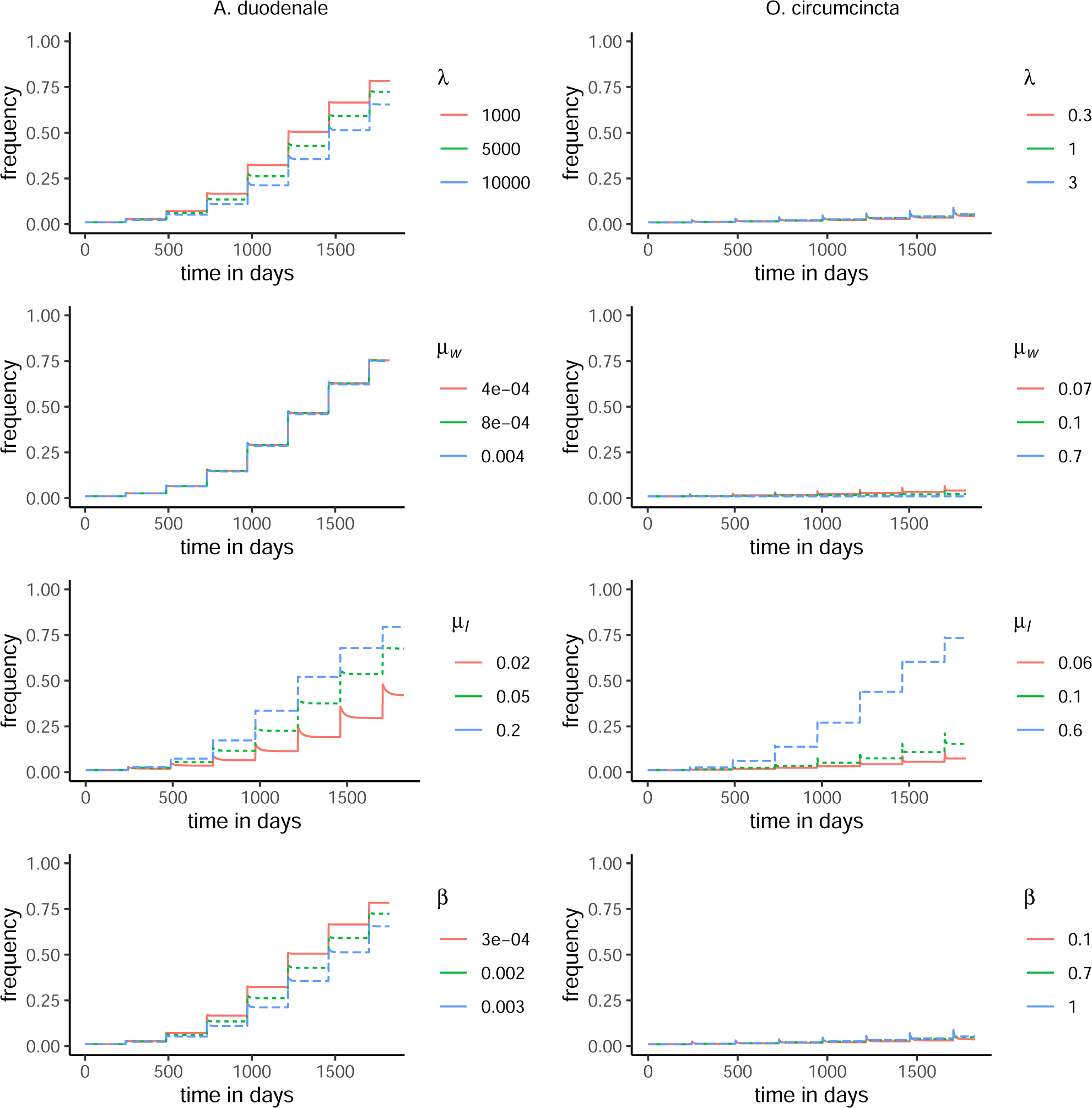
Evolutionary trajectories of *x*(*t*) with varying parameter values for fixed treatment frequency (1*/τ* = 1.5) and coverage (*c* = 1). All parameters except one (indicated in each panel) are fixed at values corresponding to *A. duodenale* and *O. circumcincta* are given in Table 1. Genotype-dependent drug efficacies are fixed to be *µ_AA_* = 0.1*, µ_AB_* = 0.5*, µ_BB_* = 0.9.

We find that increases in the larval death rate *µ_w_* increase the rate of evolution by indirectly increasing the worm to larvae ratio, Figure 8. On the other hand, increases in the adult worm death rate decrease the rate of evolution by indirectly decreasing the worm to larvae ratio. Similarly, increases in the contact rate, *β* or the egg production, *λ*, decrease the rate of evolution (Figure 8).

#### 4.3.2 Treatment Regime

Finally, we asked how different treatment strategies quantitatively impact the worm population dynamics and resistance evolution. We compared the dynamics over a range of (a) treatment frequencies 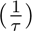 with coverage fixed, (b) coverage (*c*) with treatment frequency fixed, and (c) both simultaneously, keeping τ fixed. The last scenario allows us to compare across treatment strategies so the total number of people treated over a longer time span is constant.

We find that treating more often or treating with higher coverage both speed up the evolution of resistance. This is because selection events are more frequent and stronger, respectively. Despite this increased rate of resistance evolution, we find that over time the total number of worms within the host population is always smaller with either increased coverage or treatment frequency (not shown).

Varying both *c* and *τ* simultaneously, while 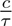 is constant, we ask is it better to treat more often with less coverage or less often with higher coverage? Figure 9 shows results for *A. duodenale*. In the absence of resistance evolution, treating less often with higher coverage (blue curve in Figure 9) reduces the number of worms within the hosts more effectively than treating more often with less coverage (red curve in Figure 9). With resistance evolution, we find that treating less often at higher coverage initially reduces the number of worms in the hosts more effectively. However, after some time period, there is evolutionary rescue and the worm burden in hosts increases more rapidly, which in our example happens after roughly 15 years. This is because in this treatment regime the evolution of resistance is faster.

**Figure 9:**
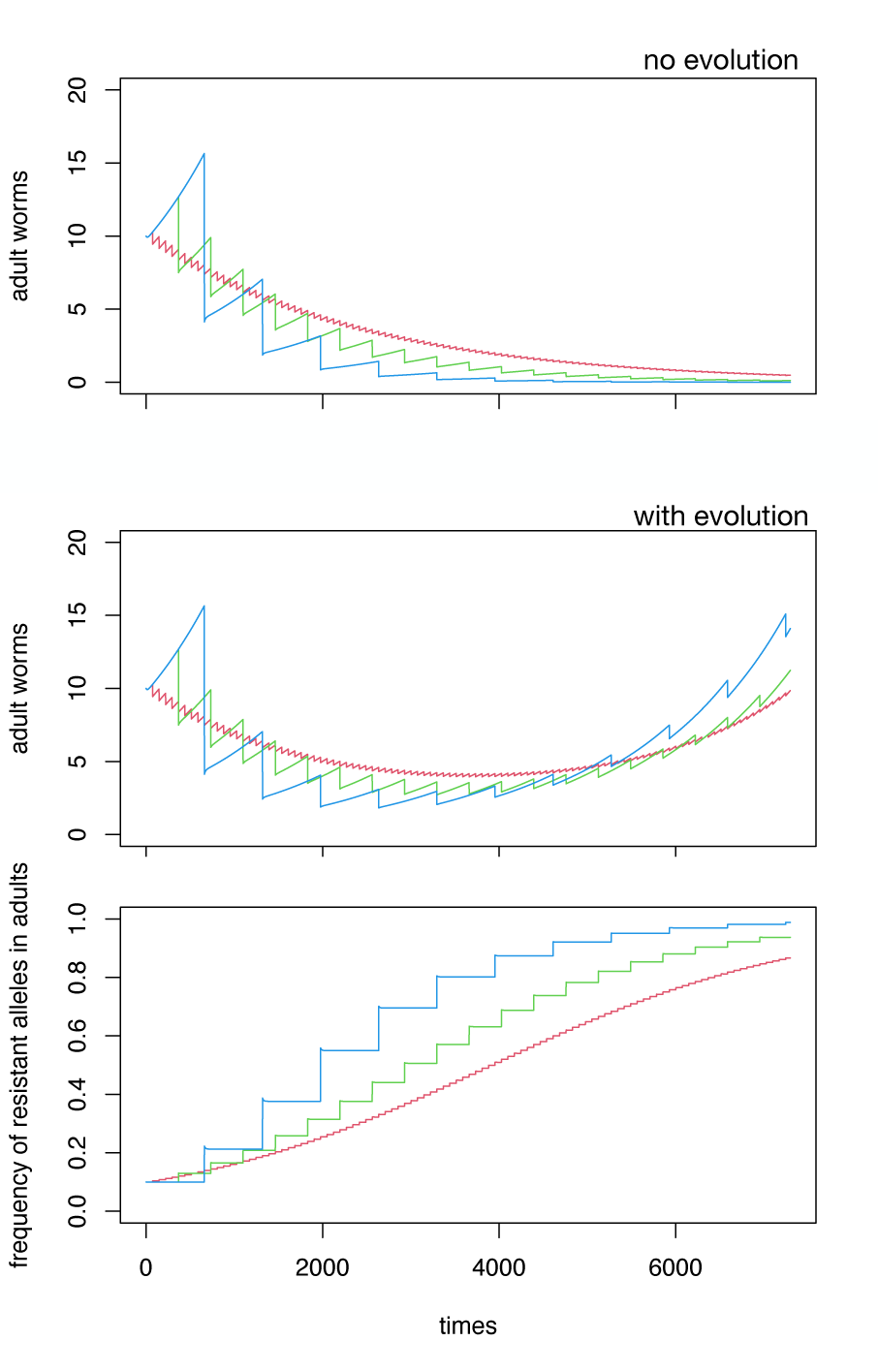
Effect of frequency versus coverage. Each panel shows three treatment regimes: low coverage (*c* = 0.1) at a high frequency (1*/τ* = 5) is in red, medium coverage (*c* = 0.5) and frequency (1*/τ* = 1) is in green and high coverage (*c* = 0.9) and low frequency (1*/τ* = 0.555) is in blue. Note that *c/τ* is held constant at 0.5 for each regime. (A) the number of adult worms in the host population over time with no resistance evolution. (B-C) number of adult worms and frequency of resistant alleles with resistance evolution. All other parameters are from table for *A. duodenale*.

## 5 Discussion

In this work, we developed a mechanistic understanding of how fundamental aspects of parasite and host biology impact the population and resistance evolution dynamics of STH. We focused on a simple model that ignores many details included in more realistic models of soil-transmitted helminth transmission and treatment. We give an explicit derivation of our model, which includes two life cycle stages and a genetic structure that only impacts the drug efficacy. The reason was to allow for the development of rigorous analytically tractable results and to focus on a mechanistic understanding of how fundamental aspects of the basic biology of these organisms impact the dynamics of resistance evolution in parasites. Generally, the human-infecting species have a smaller host-reservoir contact rate, adult worm death rate and worm to larvae ratio, but larger egg production rate than the livestock-infecting species. The livestock-infecting species have greater *R*_0_ values. Our work highlights how each species can have different propensities for persistence despite repeated frequent treatment and different propensities for resistance evolution.

Due to the simplicity of our model, we are able to characterize the qualitative dynamics with an explicit calculation of the critical treatment frequency needed for elimination, depending on the drug efficacy. Human and livestock-infecting species differ in their ecological dynamics. Generally, livestock species require much more frequent treatment to control the worm populations, which may be due to the greater *R*_0_ value. This aligns with what has been done in practice: prior to the problem of resistance, livestock farmers used to treat frequently. For example, it was common to drench lambs as often as monthly [43]. By comparing across the human-infecting species, which all have the same *R*_0_ value in our parameterizations, we observe that the treatment frequency needed for elimination depends on the parasite life cycle, even for a given *R*_0_. In other words, even in the linear model without resistance evolution, the critical treatment frequency does not depend on *R*_0_ alone.

The shape of the critical frequency curve with respect to drug efficacy provides insight into which species are most likely to evolve resistance. Given our model, our results estimate how much treatment frequency must be increased to continue on the path towards elimination in the face of resistance evolution. In Figure 2, the steeper the critical frequency curve, the greater the impact small changes in overall drug efficacy can have on the parasite dynamics. In our model, resistance that leads to small changes in drug efficacy has the most pronounced effect for *A. duodenale* and *H. contortus*, in human- and livestock-infecting species, respectively.

Additionally, in our model, most human-infecting species evolve resistance faster than livestock-infecting species. This is surprising for two reasons. One, the worm to larvae ratio, which speeds up resistance evolution, is much larger in the livestock-infecting species than the human-infecting species. However, this is countered by the high contact rate and lower fecundity of the livestock-infecting species. Both factors have a direct effect to slow down evolution, for a given worm to larvae ratio. The second reason why it is surprising is empirical: resistance has been detected to a much lesser degree in humans than in livestock populations. This could be due to the history of lower treatment frequency and coverage in humans. It may also be due in part to differences in the number and methods of empirical studies aimed at detecting resistance.

When testing the effects of treatment frequency (and coverage) on the dynamics, we expected to observe that there was an optimal treatment frequency (and coverage) to cause the parasite population to decline, since on the one hand each treatment event results in increased mortality of the parasites but on the other it leads to evolution that reduces their mortality. What we found was that increasing treatment frequency (when coverage is fixed) always caused an overall more pronounced decline in the parasite dynamics. Hence, it was better to treat more often to control the population, in our model, despite the faster evolution to resistance. This is a consequence of our model being linear and that there were no evolutionary fitness costs of resistance (which has been supported from an empirical standpoint at least for benzimidazole drugs [44]–[46].

Additionally, we tested the impact of treatment strategies when the intensity of treatment, 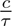 was fixed, but varying coverage and frequency. Although we can’t recommend specific treatment solutions because of our simplified model, we found that in the short term, regimes with higher coverage and lower frequency led to a larger reduction in the worm population compared to those with lower coverage and higher frequency, for a given intensity of treatment 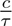. However, in the long term, the higher coverage regime resulted in faster evolution of resistance and a faster rebound.

This model and analyis framework are intended to be a baseline for further investigation, including understanding the impact of additional more complex features of soil-transmitted helminth biology on the evolution of resistance. In particular, it is well accepted that density-dependent fecundity, mortality and mating, are important to the ecological dynamics of these parasites [33], [47]. Moreover, these are amplified when considering heterogeneity in the distribution of worms amongst hosts [48]. An important paper on anthelmintic resistance [33] incorporated three different forms of density dependence into a transmission model to examine how different forms of density dependence affect the rate of the evolution of resistance. They found that incorporating density-dependent fecundity was associated with a faster evolution of resistance than density-dependent establishment into hosts and density-dependent mortality. It will be worthwhile to examine how these types of density-dependencies affect the optimal strategies for treating to minimize worm burden.

From an evolutionary perspective, in this work, we assumed a single genetic locus confers resistance and that there was no fitness cost associated with the resistance allele. That is, all genetic types were equivalent except in their drug efficacy. Experimental studies in model organisms, such as *C. elegans*, aim to evaluate whether there are any costs to resistance genes but there are still many unknowns [46]. Theoretical studies that explore the implications of costs to resistance on its evolution can motivate or guide experimental work. Conversely, empirical work to determine the genetic basis and associated costs of resistance can inform better models. Indeed, recent work uses simulations of a newly developed individual-based model shows that the rate of the resistance evolution depends on the whether the trait is monogenic or polygenic [49]. The genetic basis of these traits may be more complex than we typically assume in our models. Hence, uncovering and then incorporating this reality into the model may provide more insight into our understanding of when and how resistance evolves.

## 6 Supplements

**6.1 Model Description**

**6.2 Parameterizations and Table**

**6.3 Critical Time Analysis**

**6.4 Additional Figures**

## Supporting information

SI1 Model Derivation

SI2 Parameterizations

SI3 Critical Time Analysis

SI4 Mathematica Notebook

## Acknowledgements

This project has been made possible with support of the Oregon State University College of Science Research and Innovation Seed (SciRIS) program (https://beav.es/ihi) from the Disease Mechanism and Prevention Fund to SP, the National Science Foundation Postdoctoral Research Fellowship in Biology (2208947) to KL and was the National Science Foundation Grant (DMS-1951759) and the Simons Foundation/SFARI (638193) to HG.

## Notes

https://github.com/swapatel/STH\_baseline\_eco-evo

### Competing Interest Statement

The authors have declared no competing interest.

