## Supplementary material for "Eco-evolutionary dynamics of anthelmintic resistance in soil-transmitted helminths": SI1 Model Derivation

### Supplementary Information I: Model Derivation

November 13, 2023

In this supplement, we derive the model, with particular emphasis on the evolutionary dynamics. We begin with the dynamics in between treatment pulses and then derive the expressions for the instantaneous effects of pulsed treatments.

#### 1 Dynamics in Between Treatment Pulses

Soil-transmitted helminths spend part of their life cycle as adults in a host and as larvae in a soil environment. We assume that there is a natural death of the adult worms, $\mu_w$ and larvae, $\mu_l$. Additionally, adult worms can die if their host dies, at rate $\mu_h$. Female adult worms produce eggs that get secreted into the environment and turn into larvae at a rate $\lambda$ and hosts take in larvae at rate $\beta N$, where $N$ is a constant number of hosts.

We assume there is a single locus in the genome of the parasite with two possible alleles and that the genotype of an individual determines its response to the treatment drug, i.e., the drug efficacy. We assume that one allele, labeled "A", is a resistant allele, while the other possible alelle, labeled "B", is a susceptble allele. Let $x(t)$ and $y(t)$ be the frequency of the A allele in the hosts and soil environment, respectively.

As these parasites are diploid, there three possible genotypes: $AA, AB, BB$. All life history parameters are independent of the genotype, except for at treatment pulses. Hence, if we let $W_{ij}$ and $L_{ij}$ be the density of adult worms and larvae, respectively, with genotype $ij$, we express the dynamics of the adult worms in this genotype class as

$$\frac{dW_{ij}}{dt} = \beta N L_{ij} - (\mu_w + \mu_h) W_{ij} \tag{1}$$

Larvae of type $ij$ are produced from sexual mating between two worms, one with an $i$ allele and the other with $j$ allele. The dynamics of the larvae of this genotype class are

$$\frac{dL_{ij}}{dt} = \ell_{ij}(W_{AA}, W_{AB}, W_{BB}) - (\mu_l + \beta N) L_{ij} \tag{2}$$

where $\ell_{ij}$ is the rate at which larvae of type $ij$ are produced and is a function of the adult worms of each type. We assume that half of the adult worm population is female, each female produces eggs at a rate $\lambda$ independent of genotype and that these eggs are randomly fertilized by a male adult worm. Bookkeeping the possibilities, then

$$\ell_{AA} = \frac{\lambda}{2}\big[W_{AA}(\frac{2W_{AA}+W_{AB}}{2W}) + \frac{1}{2}W_{AB}(\frac{2W_{AA}+W_{AB}}{2W})\big] \tag{3a}$$

$$\ell_{AB} = \frac{\lambda}{2}\big[W_{AA}(\frac{2W_{BB}+W_{AB}}{2W}) + W_{AB}(\frac{W_{AA}+W_{AB}+W_{22}}{2W}) + W_{BB}(\frac{2W_{AA}+W_{AB}}{2W})\big] \tag{3b}$$

$$\ell_{BB} = \frac{\lambda}{2}\big[W_{BB}(\frac{2W_{BB}+W_{AB}}{2W}) + \frac{1}{2}W_{AB}(\frac{2W_{BB}+W_{AB}}{2W})\big] \tag{3c}$$

Here $W = W_{AA} + W_{AB} + W_{BB}$ is the total adult worm population. Similarly, let $L = ...$ be the total larvae in the soil.

Then,

$$x = \frac{2W_{AA}+W_{AB}}{2W} \tag{4}$$

and

$$y = \frac{2L_{AA}+L_{AB}}{2L} \tag{5}$$

With this, equations 3 can be more simply expressed as

$$\ell_{AA} = \frac{\lambda}{2}\big[W_{AA}x + \frac{1}{2}W_{AB}x\big] \tag{6a}$$

$$\ell_{AB} = \frac{\lambda}{2}\big[W_{AA}(1-x) + \frac{1}{2}W_{AB} + W_{BB}x\big] \tag{6b}$$

$$\ell_{BB} = \frac{\lambda}{2}\big[W_{BB}(1-x) + \frac{1}{2}W_{AB}(1-x)\big] \tag{6c}$$

and we can express the dynamics of the total worm and larvae populations as

$$\frac{dW}{dt} = \beta NL - (\mu_w + \mu_h)W \tag{7}$$

$$\frac{dL}{dt} = \frac{\lambda}{2}W - (\mu_l + \beta N)L \tag{8}$$

Furthermore, using quotient rule on 4 and 5 and applying 1 and 2, we arrive at

$$\frac{dx}{dt} = \beta N\big[\frac{L}{W}(y-x)\big] \tag{9}$$

$$\frac{dy}{dt} = \frac{\lambda}{2}\big[\frac{W}{L}(x-y)\big] \tag{10}$$

### 2 Dynamics at Pulsed Treatment Times

We let $\mu_{ij}$ be the drug efficacy, i.e., the probability of dying at the time of treatment, for individuals with genotype $ij$. We let $c$ be the coverage, i.e., the proportion of the total host population that is treated and $\tau$ be the time in between pulsed treatments. Then, for $n \in N$,

$$W_{ij}(n\tau^+) = (1 - c\mu_{ij})W_{ij} \tag{11}$$

is the number of adult worms of genotype $ij$ after treatment.

Assuming Hardy-Weinberg equilibrium, we have that

$$W_{AA} = x^2 W \tag{12a}$$
$$W_{AB} = 2x(1-x)W \tag{12b}$$
$$W_{BB} = (1-x)^2 W \tag{12c}$$

Under this assumption, we express the pulse effect on the whole adult worm population as

$$W(n\tau^+) = W_{AB}(n\tau^+) + W_{AB}(n\tau^+) + W_{BB}(n\tau^+) \tag{13}$$
$$= (1 - c\overline{\mu})W \tag{14}$$

where $\overline{\mu} = x^2\mu_{AA} + 2x(1-x)\mu_{AB} + (1-x)^2\mu_{BB}$ is the average drug efficacy.

Similarly, the pulse effects on the frequency of the resistant allele (from selection) is

$$x(n\tau^+) = \frac{2W_{AA}(n\tau^+) + W_{AB}(n\tau^+)}{2W(n\tau^+)} \tag{15}$$
$$= \frac{1 - c\overline{\mu_A}}{1 - c\overline{\mu}} x \tag{16}$$

where $\overline{\mu_A} = x\mu_{AA} + (1-x)\mu_{AB}$ is the marginal drug efficacy for an allele of type A. One can interpret this from the perspective of allele A as its drug efficacy. This is the probability of one allele A being in a genotype AA or AB times the respective drug efficacy of that genotype. Note that since $x$ is dynamic in time, so are $\overline{\mu}$ and $\overline{\mu_A}$

Altogether, equations 7-10, 14, and 16 are the model.
