## Supplementary material for "Eco-evolutionary dynamics of anthelmintic resistance in soil-transmitted helminths": SI2 Parameterizations

### Supplementary Information: Parameterization

November 7, 2023

We parameterized the model for human-infecting soil-transmitted helminth (STH) species (Main Text Table 1) using life history parameters [1]–[3], for livestock-infecting species species (Main Text Table 2) using life history parameters from [4]. For other parameters, we used information on populations, as described in more detail below. All rate parameters have been converted to daily rates.

### 1 Human-infecting species

We consider three human-infecting STH species: *A. lumbricoides, A. duodenale, T. trichuris.*

#### 1.1 Parasite Biology Parameters

The adult worm death rates $\mu_w$ and the egg / larvae death rates $\mu_l$ for *A. lumbricoides* and *A. duodenale* were taken from [3] (Imperial College Model) and for *T. trichuris* [2]. The egg production rates $\lambda$ for all worm species were the lower values of the range given in [2] converted from a per day to a per year rate.

#### 1.2 Host Parameters

Human life span was assumed to be 60 years, which we converted into the death rate per day, $\mu_h$. 60 years is shorter than modern life expectancies but corresponds to about two human generations. The host life expectancy is much longer than the adult worm life expectancy and therefore has a negligible impact on modelled outcomes.For human host density, $N$, we used the estimated population density from Himachal Pradesh, India (Population Census India 2011, `https//www.census2011.co.in`). The contact rates $\beta$ for all human parasite species were calculated using the following expression for $R_0$ and assuming an $R_0$ value of 1.5

$$R_0 = \frac{\frac{1}{2}\lambda\beta N}{(\mu_h + \mu_a)(\mu_l + \beta N)}$$

An $R_0$ value of 1.5 lies within the typically observed range for human soil-transmitted helminth parasites in low- to moderate-endemicity settings [2].

### 2 Livestock-infecting species

We consider three livestock-infecting STH species and use parameter values from [4]: *T. colubriformes* (parameters from Australia), *O. circumcincta* (parameters from UK), *H. contortus* (parameters from Australia).

#### 2.1 Parasite Biology Parameters

Previous veterinary helminth parasite models include two parameters that are not routinely used in human helminth parasite models because they are difficult to measure in the human context, probability of larvae to develop into adult worms inside the host $p$ and the probability of eggs to develop into infective larvae $q$. To make the models comparable, we calculate the effective contact rate with the environmental reservoir of infectious eggs and larvae $\beta_{eff}$ and the effective egg production rate per female worm $\lambda_{eff}$ for the veterinary helminth parasite model using the values from [4]:

$$\beta_{eff} = p\beta \tag{1}$$

$$\lambda_{eff} = q\lambda \tag{2}$$

Thus, we implicitly assume $q = 1$ and $p = 1$ in the human helminth parasite model. Note that $\beta$ occurs twice in the ecological helminth model, once in the term describing human infections and once in the term describing depletion of the environmental reservoir by egg / larval death and uptake of eggs / larvae by host animals. Strictly, the value of $\beta_{eff}$ is only correct for the infection term, but since uptake of eggs / larvae by host animals is small relative to the egg / larval death rate, we assume that changes the results from this inaccuracy are negligible.

In seasonal settings, the probability that larvae are taken up by a host animal and develop into adult worms varies in different months of the year [4]. Seasonal variation in infection probability has also been observed for human helminth parasites [5]. But it is less pronounced and therefore commonly ignored in helminth infection models. For simplicity, we ignore seasonal effects on the infection rate in both human and veterinary parasites. For veterinary parasites, we use the annual mean of the infectious contact probability as given in [4] in our calculation of the effective contact rate $\beta_{eff}$.

The adult worm death rate $\mu_w$ in veterinary helminth parasites is affected by the host immune response [4]. However, human hosts do not develop an effective immune response against soil-transmitted helminth infection. For comparability between veterinary and human parasite models, we ignore the effect of host immunity on adult worm death rate.

Reported larval death rates $\mu_l$ for veterinary helminth parasites vary by environmental setting [4]. In our analysis we use the values reported for Australia or the UK depending on helminth species, as indicated above.

#### 2.2 Host Parameters

The host population density $N$ is measured in hosts per $m^2$ in the veterinary helminth parasite model and lies in the range of $10^{-3}$ [4]. We assume a sheep life expectancy of 10 years [6].

Previous veterinary helminth parasite models also include immunity, but models for soil-transmitted helminth infections in humans do not. To be able to compare all other parameters between veterinary and human helminth parasites, we ignore immunity in the veterinary helminth parasite models.
