## Supplementary material for "Eco-evolutionary dynamics of anthelmintic resistance in soil-transmitted helminths": SI3 Critical Time Analysis

### Supplementary Information: Critical Time Analysis

November 7, 2023

In this document, we determine an upper bound on the time between treatments required to eliminate the parasites from the host population in terms of parameters of our model, assuming there is no evolution (fixed drug efficacy). We show that this critical time is the solution to a transcendental equation. In a separate supplemental Mathematica notebook, we numerically solve this transcendental equation for the critical time for particular species.

#### 1 Model

Let $W$ be the adult worm burden in the host population and $L$ the larvae in the soil environment. The model for a fixed drug efficacy (without any evolution) we study is

$$\begin{aligned} \frac{dW}{dt} &= \beta N L - (\mu_h + \mu_a) W && t \neq n\tau \\ \frac{dL}{dt} &= \lambda W - (\mu_l + \beta N) L && t \neq n\tau \\ W(n\tau^+) &= (1 - c\mu) W && t = n\tau \end{aligned} \tag{1}$$

with positive parameters $\beta, N, \lambda, \mu_h, \mu_w, \mu_l, \mu$. Here, $\tau$ is the time between periodic drug treatments with $n$ in the natural numbers. Additionally, $c$ is the coverage (the proportion of the population treated) and $\mu$ is the drug efficacy, which has the effect of reducing some proportion of the total worm population at the exact time of treatment.

Notice that, in the absence of treatment, this is a linear model whose dynamics can be captured by a matrix

$$A = \begin{bmatrix} -(\mu_h + \mu_a) & \beta N \\ \lambda & (\mu_l + \beta N) \end{bmatrix}$$

It follows from the Perron-Frobenius theorem, that $A$ has two real eigenvalues $\lambda_1 > \lambda_2$. Moreover, $A$ has exactly one eigenvector $\vec{v_1}$ in the positive quadrant, and $\vec{v_1}$ is the eigenvector corresponding to eigenvalue $\lambda_1$. Then, we have that $\lambda_1$ is positive if and only if

$$(\mu_h + \mu_a)(\mu_l + \beta N) < \lambda \beta N \tag{2}$$

In this case, we have exponential growth of the parasite without treatment. We will assume this condition is met, because otherwise, the parasite goes extinct on its own.

### 2 Critical Time

We ask how often must treatment need to be applied to prevent growth and drive the parasite to extinction? We find a critical time $\tau^*$ such that for $\tau > \tau^*$, we have that $W(t), L(t) \to \infty$ and for $\tau < \tau^*$, we have that $W(t), L(t) \to 0$. The existence and uniqueness of such a critical time follows from Theorem 2 in [1].

#### 2.1 Solutions without the treatment pulse

Since we have a linear system in the absence of treatment, we have explicit solutions. For initial condition $W_0 = W(0), L_0 = L(0)$, solutions are

$$\begin{bmatrix} W(t) \\ L(t) \end{bmatrix} = r_1 e^{\lambda_1 t} \vec{v_1} + r_2 e^{\lambda_2 t} \vec{v_2} \tag{3}$$

where $\lambda_1, \lambda_2$ are eigenvalues (both real), with corresponding eigenvectors

$$\vec{v_1} = \begin{bmatrix} a \\ 1 \end{bmatrix}, \vec{v_2} = \begin{bmatrix} b \\ 1 \end{bmatrix} \tag{4}$$

and

$$\begin{bmatrix} r_1 \\ r_2 \end{bmatrix} = \begin{bmatrix} \vec{v_1} \vec{v_2} \end{bmatrix}^{-1} \begin{bmatrix} W_0 \\ L_0 \end{bmatrix}. \tag{5}$$

#### 2.2 Solutions with the treatment pulse

For model 1 and fixed $\tau > 0$, we can define a discrete time map for the parasite and larvae right after treatment. Let $W_{n\tau}$ and $L_{n\tau}$ be the density of adult worms and larvae at the time point right after the $n^{th}$ treatment. Then,

$$\begin{aligned} W_{(n+1)\tau} &= (1 - c\mu)[r_1 a e^{\lambda_1 \tau} + r_2 b e^{\lambda_2 \tau}] \\ L_{(n+1)\tau} &= r_1 e^{\lambda_1 \tau} + r_2 e^{\lambda_2 \tau} \end{aligned} \tag{6}$$

where

$$\begin{bmatrix} r_1 \\ r_2 \end{bmatrix} = \begin{bmatrix} \vec{v_1} \vec{v_2} \end{bmatrix}^{-1} \begin{bmatrix} W_{n\tau} \\ L_{n\tau} \end{bmatrix}.$$

with $\vec{v_1}$ and $\vec{v_2}$ defined as in 2.1.

A non-zero fixed point $(\hat{W}, \hat{L})$ of this discrete time map satisfies

$$\begin{aligned} \hat{W} &= a r_1 + b r_2 = (1 - c\mu)[a r_1 e^{\lambda_1 \tau} + b r_2 e^{\lambda_2 \tau}] \\ \hat{L} &= r_1 + r_2 = r_1 e^{\lambda_1 \tau} + r_2 e^{\lambda_2 \tau} \end{aligned} \tag{7}$$

which can only have a solution if $\tau = \tau^*$, given by

$$\frac{e^{\lambda_2 \tau^*} - 1}{1 - e^{\lambda_1 \tau^*}} = \frac{b(\rho e^{\lambda_2 \tau^*} - 1)}{a(1 - \rho e^{\lambda_1 \tau^*})} \tag{8}$$

where $\rho = 1 - c\mu$. This is a transcendental equation. Hence, we numerically find the critical $\tau^*$ in Mathematica for different species. (See Mathematica notebook.)
