## Supplementary material for "Eco-evolutionary dynamics of anthelmintic resistance in soil-transmitted helminths": SI4 Mathematica Notebook

### Critical Time Analysis

In the first section of this notebook, we determine the critical time between treatments needed for periodic cycling. We do this for different species (with different parameters) of human-infecting and vet-infecting soil-transmitted helminths. In the second section “Categorizing Behaviors”, we determine the qualitative behavior of parameter sets inputted from a csv file.

Model equations

In[∘]:=
```
fW = β * Nhost * L - (μh + μa) * W;
fL = 0.5 * W * λ - (μl + β * Nhost) * L;
```

Eigenvalues, eigenvectors

In[∘]:=
```
Jacob = Simplify[D[{fW, fL}, {{W, L}}]];
```

In[∘]:=
```
eigensys = Eigensystem[Jacob]
λ1 = eigensys[[1]][[1]]
λ2 = eigensys[[1]][[2]]
```

Out[∘]=
$$\{\{0.5\,(-1.\,\mathrm{Nhost}\,\beta - 1.\,\mu a - 1.\,\mu h - 1.\,\mu l + 1.\,\sqrt{((1.\,\mathrm{Nhost}\,\beta + 1.\,\mu a + 1.\,\mu h + 1.\,\mu l)^2 - 4.\,(-0.5\,\mathrm{Nhost}\,\beta\,\lambda + 1.\,\mathrm{Nhost}\,\beta\,\mu a + 1.\,\mathrm{Nhost}\,\beta\,\mu h + 1.\,\mu a\,\mu l + 1.\,\mu h\,\mu l))}),$$
$$-0.5\,(1.\,\mathrm{Nhost}\,\beta + 1.\,\mu a + 1.\,\mu h + 1.\,\mu l + 1.\,\sqrt{((1.\,\mathrm{Nhost}\,\beta + 1.\,\mu a + 1.\,\mu h + 1.\,\mu l)^2 - 4.\,(-0.5\,\mathrm{Nhost}\,\beta\,\lambda + 1.\,\mathrm{Nhost}\,\beta\,\mu a + 1.\,\mathrm{Nhost}\,\beta\,\mu h + 1.\,\mu a\,\mu l + 1.\,\mu h\,\mu l))})\},$$
$$\{\{\frac{1}{\lambda}1.\,(1.\,\mathrm{Nhost}\,\beta - 1.\,\mu a - 1.\,\mu h + 1.\,\mu l + 1.\,\sqrt{((1.\,\mathrm{Nhost}\,\beta + 1.\,\mu a + 1.\,\mu h + 1.\,\mu l)^2 - 4.\,(-0.5\,\mathrm{Nhost}\,\beta\,\lambda + 1.\,\mathrm{Nhost}\,\beta\,\mu a + 1.\,\mathrm{Nhost}\,\beta\,\mu h + 1.\,\mu a\,\mu l + 1.\,\mu h\,\mu l))}),\ 1.\},$$
$$\{-\frac{1}{\lambda}1.\,(-1.\,\mathrm{Nhost}\,\beta + 1.\,\mu a + 1.\,\mu h - 1.\,\mu l + 1.\,\sqrt{((1.\,\mathrm{Nhost}\,\beta + 1.\,\mu a + 1.\,\mu h + 1.\,\mu l)^2 - 4.\,(-0.5\,\mathrm{Nhost}\,\beta\,\lambda + 1.\,\mathrm{Nhost}\,\beta\,\mu a + 1.\,\mathrm{Nhost}\,\beta\,\mu h + 1.\,\mu a\,\mu l + 1.\,\mu h\,\mu l))}),\ 1.\}\}\}$$

Out[∘]=
$$0.5\,(-1.\,\mathrm{Nhost}\,\beta - 1.\,\mu a - 1.\,\mu h - 1.\,\mu l + 1.\,\sqrt{((1.\,\mathrm{Nhost}\,\beta + 1.\,\mu a + 1.\,\mu h + 1.\,\mu l)^2 - 4.\,(-0.5\,\mathrm{Nhost}\,\beta\,\lambda + 1.\,\mathrm{Nhost}\,\beta\,\mu a + 1.\,\mathrm{Nhost}\,\beta\,\mu h + 1.\,\mu a\,\mu l + 1.\,\mu h\,\mu l))})$$

Out[∘]=
$$-0.5\,(1.\,\mathrm{Nhost}\,\beta + 1.\,\mu a + 1.\,\mu h + 1.\,\mu l + 1.\,\sqrt{((1.\,\mathrm{Nhost}\,\beta + 1.\,\mu a + 1.\,\mu h + 1.\,\mu l)^2 - 4.\,(-0.5\,\mathrm{Nhost}\,\beta\,\lambda + 1.\,\mathrm{Nhost}\,\beta\,\mu a + 1.\,\mathrm{Nhost}\,\beta\,\mu h + 1.\,\mu a\,\mu l + 1.\,\mu h\,\mu l))})$$

In[•]:=
```
eigenvec = Eigenvectors[Jacob];
inveigenvec = Inverse[Transpose[eigenvec]];
c1c2 = inveigenvec.{W0, L0};
c1 = c1c2[[1]];
c2 = c1c2[[2]];
```

#### Solution to critical equation

A critical frequency of MDA exists only if without MDA, the worm population blows up (i.e., requires a negative determinant of the Jacobian).

Ascaris

In[•]:=
```
IC = {W0 → 100, L0 → 3000};
ParmsAsc = {β → 0.04 / 365, Nhost → 0.000123,
   μh → 0.0167 / 365, μa → 1 / 365, λ → 3.65 * 10^6 / 365, μl → 6 / 365};
Tr[Jacob] /. ParmsAsc
Det[Jacob] /. ParmsAsc
λ1 /. ParmsAsc
λ2 /. ParmsAsc
eigenvec /. ParmsAsc
```

Out[•]= -0.0192238

Out[•]= -0.0000216085

Out[•]= 0.00106504

Out[•]= -0.0202889

Out[•]= $\{\{3.50068\times10^{-6}, 1.\}, \{-7.70104\times10^{-7}, 1.\}\}$

Hookworm

In[•]:=
```
IC = {W0 → 100, L0 → 3000};
ParmsHk = {β → 0.35 / 365, Nhost → 0.000123,
   μh → 0.0167 / 365, μa → 0.5 / 365, λ → 1.095 * 10^6 / 365, μl → 30 / 365};
Tr[Jacob] /. ParmsHk
Det[Jacob] /. ParmsHk
eigenvec /. ParmsHk
```

Out[•]= -0.0836075

Out[•]= -0.0000605656

Out[•]= $\{\{0.0000552734, 1.\}, \{-1.42257\times10^{-6}, 1.\}\}$

Trichuris

In[∘]:=
```
IC = {W0 → 100, L0 → 3000};
ParmsTri = {β → 0.62 / 365, Nhost → 0.000123,
   μh → 0.0167 / 365, μa → 1 / 365, λ → 7.3 * 10 ^ 5 / 365, μl → 18.25 / 365};
Tr[Jacob] /. ParmsTri
Det[Jacob] /. ParmsTri
eigenvec /. ParmsTri
```

Out[∘]= -0.0527857

Out[∘]= -0.000069657

Out[∘]= $\{\{0.0000512884, 1.\}, \{-4.07366 \times 10^{-6}, 1.\}\}$

TcolubNZ

In[∘]:=
```
IC = {W0 → 100, L0 → 3000};
ParmsTc = {β → 511 / 365, Nhost → 0.0015,
   μh → 0.1 / 365, μa → 17.89 / 365, λ → 3285 / 365, μl → 8.4 / 365};
Tr[Jacob] /. ParmsTc
Det[Jacob] /. ParmsTc
λ1 /. ParmsTc
λ2 /. ParmsTc
eigenvec /. ParmsTc
```

Out[∘]= -0.0744014

Out[∘]= -0.0082122

Out[∘]= 0.060759

Out[∘]= -0.13516

Out[∘]= {{0.0190828, 1.}, {-0.0244548, 1.}}

TcolubAUS

```
In[•]:= IC = {W0 → 100, L0 → 3000};
ParmsTcA = {β → 1124.2 / 365, Nhost → 0.001,
    μh → 0.1 / 365, μa → 17.89 / 365, λ → 2463.75 / 365, μl → 28.4 / 365};
Tr[Jacob] /. ParmsTcA
Det[Jacob] /. ParmsTcA
λ1 /. ParmsTcA
λ2 /. ParmsTcA
eigenvec /. ParmsTcA

Out[•]= -0.130176

Out[•]= -0.00640821

Out[•]= 0.038085

Out[•]= -0.168261

Out[•]= {{0.0352513, 1.}, {-0.0258882, 1.}}
```

OcircNZ

```
In[•]:= IC = {W0 → 100, L0 → 3000};
ParmsOc = {β → 1080.4 / 365, Nhost → 0.0015,
    μh → 0.1 / 365, μa → 40.9 / 365, λ → 4380 / 365, μl → 8.49 / 365};
Tr[Jacob] /. ParmsOc
Det[Jacob] /. ParmsOc
eigenvec /. ParmsOc

Out[•]= -0.140029

Out[•]= -0.0235285

Out[•]= {{0.0210499, 1.}, {-0.0351546, 1.}}
```

OcircUK

```
In[•]:= IC = {W0 → 100, L0 → 3000};
ParmsOcUK = {β → 956.3 / 365, Nhost → 0.002,
    μh → 0.1 / 365, μa → 14.9 / 365, λ → 1752 / 365, μl → 11.12 / 365};
Tr[Jacob] /. ParmsOcUK
Det[Jacob] /. ParmsOcUK
eigenvec /. ParmsOcUK

Out[•]= -0.0768016

Out[•]= -0.0111086

Out[•]= {{0.0456167, 1.}, {-0.0478626, 1.}}
```

HcontNZ

```
In[•]:= IC = {W0 → 100, L0 → 3000};
ParmsHc = {β → 584 / 365, Nhost → 0.0015,
   μh → 0.1 / 365, μa → 3.65 / 365, λ → 11 558.5 / 365, μl → 620.87 / 365};
Tr[Jacob] /. ParmsHc
Det[Jacob] /. ParmsHc
λ1 /. ParmsHc
λ2 /. ParmsHc
eigenvec /. ParmsHc
Out[•]= -1.71369
Out[•]= -0.0204997
Out[•]= 0.01188
Out[•]= -1.72557
Out[•]= {{0.108333, 1.}, {-0.00139918, 1.}}
```

HcontAUS

```
In[•]:= IC = {W0 → 100, L0 → 3000};
ParmsHcA = {β → 890.6 / 365, Nhost → 0.001,
   μh → 0.1 / 365, μa → 3.65 / 365, λ → 5501.8 / 365, μl → 230.36 / 365};
Tr[Jacob] /. ParmsHcA
Det[Jacob] /. ParmsHcA
eigenvec /. ParmsHcA
Out[•]= -0.643837
Out[•]= -0.0118804
Out[•]= {{0.0864455, 1.}, {-0.00374512, 1.}}
```

#### Find Critical T as a function of mu for each species

Ascaris

In[•]:=
```
ParmsAsc = {β → 0.04 / 365, Nhost → 0.000123,
    μh → 0.0167 / 365, μa → 1 / 365, λ → 3.65 * 10^6 / 365, μl → 6 / 365};
critTvec = {};
muvec = {};
For[mu = 0.1, mu ≤ .99, mu += .05,
 AppendTo[muvec, mu];
 AppendTo[critTvec, T /. NSolve[(Exp[λ2 * T] - 1) / (1 - Exp[λ1 * T]) ==
        (eigenvec[[2]][[1]] * ((1 - μ) * Exp[λ2 * T] - 1)) / (eigenvec[[1]][[1]] *
           (1 - (1 - μ) * Exp[λ1 * T])) /. μ → mu /. ParmsAsc, T, Reals][[1]]
 ]
]
critTvec
pAsc = ListLinePlot[Transpose@{muvec, critTvec},
  PlotStyle → RGBColor["#F8766D"], PlotLegends → {"A. lumbricodes"}]
```

Out[•]= {79.9298, 122.862, 167.888, 215.168, 264.923, 317.43, 373.026, 432.109, 495.154, 562.735, 635.559, 714.509, 800.712, 895.637, 1001.25, 1120.26, 1256.58, 1416.12}

Out[•]=
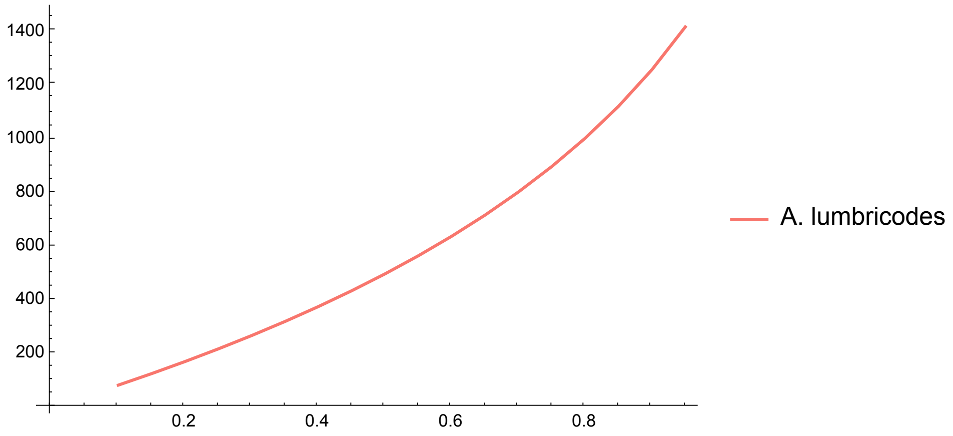


Hookworm

In[∘]:=
```
ParmsHk = {β → 0.35 / 365, Nhost → 0.000123,
   μh → 0.0167 / 365, μa → 0.5 / 365, λ → 1.095 * 10^6 / 365, μl → 30 / 365};
critTvec = {};
muvec = {};
For[mu = 0.1, mu ≤ .99, mu += .05,
 AppendTo[muvec, mu];
 AppendTo[critTvec, T /. NSolve[(Exp[λ2 * T] - 1) / (1 - Exp[λ1 * T]) ==
        (eigenvec[[2]][[1]] * ((1 - μ) * Exp[λ2 * T] - 1)) / (eigenvec[[1]][[1]] *
          (1 - (1 - μ) * Exp[λ1 * T])) /. μ → mu /. ParmsHk, T, Reals][[1]]
 ]
]
critTvec
pHk = ListLinePlot[Transpose@{muvec, critTvec},
  PlotStyle → {Dotted, RGBColor["#00BA38"]}, PlotLegends → {"A. duodenale"}]
```

Out[∘]=
```
{142.818, 220.124, 301.977, 388.945, 481.708, 581.096, 688.128, 804.078, 930.568,
 1069.71, 1224.32, 1398.26, 1597.08, 1829.1, 2107.67, 2456.27, 2922.41, 3628.21}
```

Out[∘]=

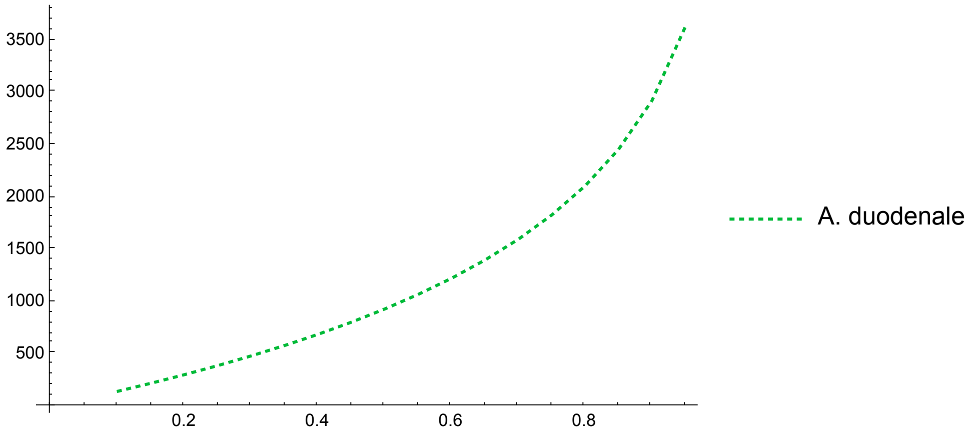


Trichuris

In[•]:=
```
ParmsTri = {β → 0.62 / 365, Nhost → 0.000123,
   μh → 0.0167 / 365, μa → 1 / 365, λ → 7.3 * 10^5 / 365, μl → 18.25 / 365};
critTvec = {};
muvec = {};
For[mu = 0.1, mu ≤ .99, mu += .05,
 AppendTo[muvec, mu];
 AppendTo[critTvec, T /. NSolve[(Exp[λ2 * T] - 1) / (1 - Exp[λ1 * T]) ==
       (eigenvec[[2]][[1]] * ((1 - μ) * Exp[λ2 * T] - 1)) / (eigenvec[[1]][[1]] *
          (1 - (1 - μ) * Exp[λ1 * T])) /. μ → mu /. ParmsTri, T, Reals][[1]]
 ]
]
critTvec
pTri = ListLinePlot[Transpose@{muvec, critTvec},
  PlotStyle → {Dashed, RGBColor["#619CFF"]}, PlotLegends → {"T. trichuris"}]
```

Out[•]=
```
{75.4591, 116.144, 159.073, 204.514, 252.78, 304.248, 359.372, 418.711, 482.965,
 553.021, 630.031, 715.532, 811.628, 921.323, 1049.11, 1202.16, 1392.99, 1646.56}
```

Out[•]=

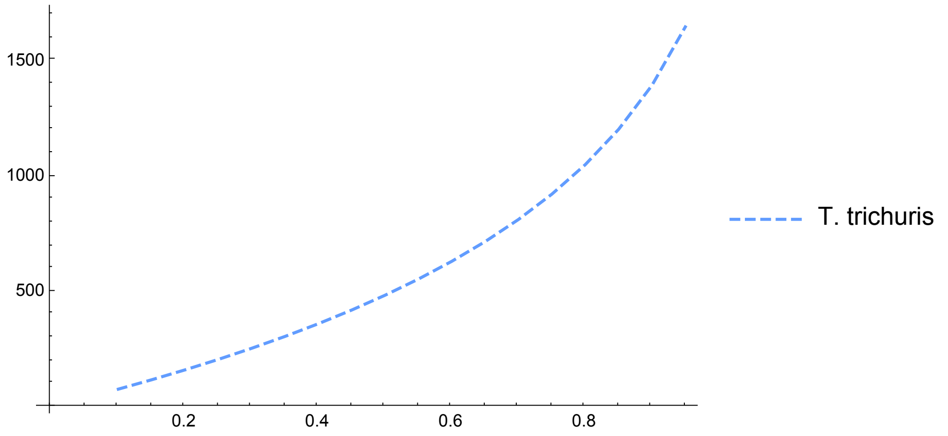


In[•]:=
```
Show[pAsc, pHk, pTri, PlotRange → {100, 3700},
 Frame -> True, FrameStyle → Directive[Black, 14]]
```

Out[•]=

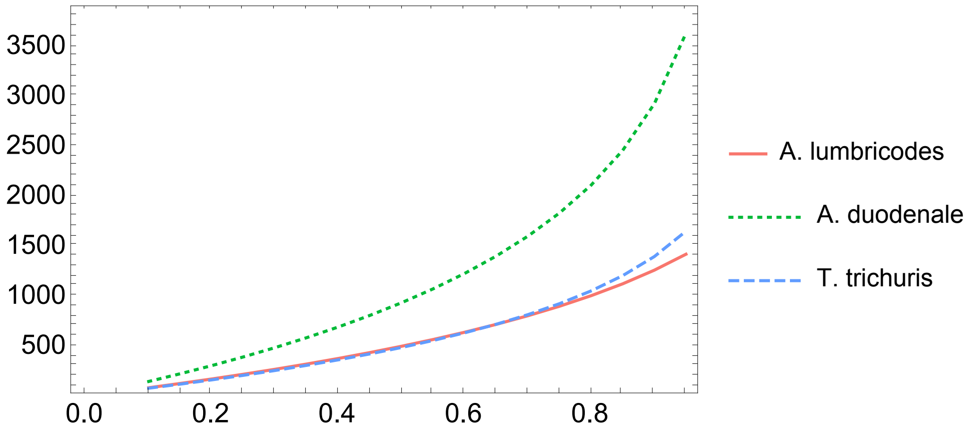

In[∘]:=
```
Show[pAsc, pHk, pTri, PlotRange → {100, 3700},
 Frame -> True, FrameStyle → Directive[Black, 14],
 PlotLegends → {{Red, {Dashed, Green}, {Dotted, Blue}},
   {"A. lumbricodes", "A. duodenale", "T. trichuris"}}]
```

Out[∘]=

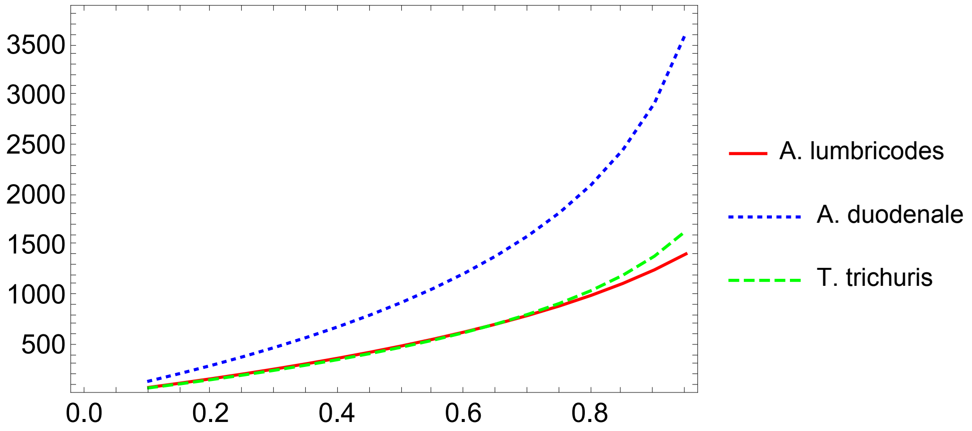


In[∘]:=
```
LineLegend[{Red, {Dashed, Green}, {Dotted, Blue}},
 {"A. lumbricodes", "A. duodenale", "T. trichuris"}]
```

Out[∘]=

A. lumbricodes
A. duodenale
T. trichuris

TColubNZ

In[∘]:=
```
ParmsTc = {β → 511 / 365, Nhost → 0.0015,
   μh → 0.1 / 365, μa → 17.89 / 365, λ → 3285 / 365, μl → 8.4 / 365};
critTvec = {};
muvec = {};
For[mu = 0.1, mu ≤ .99, mu += .05,
 AppendTo[muvec, mu];
 AppendTo[critTvec, T /. NSolve[(Exp[λ2 * T] - 1) / (1 - Exp[λ1 * T]) ==
        (eigenvec[[2]][[1]] * ((1 - μ) * Exp[λ2 * T] - 1)) / (eigenvec[[1]][[1]] *
           (1 - (1 - μ) * Exp[λ1 * T])) /. μ → mu /. ParmsTc, T, Reals][[1]]
 ]
]
critTvec
pTc = ListLinePlot[Transpose@{muvec, critTvec}, PlotStyle → RGBColor["#F564E3"]]
```

Out[∘]=
```
{0.32186, 0.495744, 0.679156, 0.872848, 1.07764, 1.29446, 1.52428, 1.76822, 2.02749,
 2.30344, 2.59754, 2.91143, 3.24691, 3.60599, 3.99084, 4.40389, 4.8478, 5.32548}
```

Out[∘]=

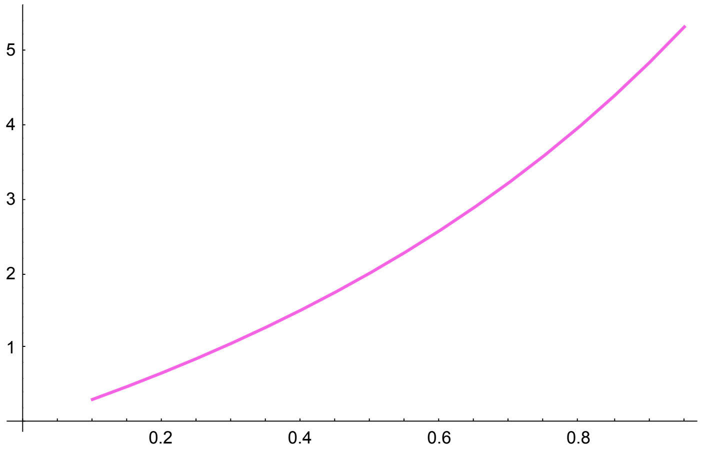


TcolubAUS

In[•]:=
```
ParmsTcA = {β → 1124.2 / 365, Nhost → 0.001,
   μh → 0.1 / 365, μa → 17.89 / 365, λ → 2463.75 / 365, μl → 28.4 / 365};
critTvec = {};
muvec = {};
For[mu = 0.1, mu ≤ .99, mu += .05,
 AppendTo[muvec, mu];
 AppendTo[critTvec, T /. NSolve[(Exp[λ2 * T] - 1) / (1 - Exp[λ1 * T]) ==
         (eigenvec[[2]][[1]] * ((1 - μ) * Exp[λ2 * T] - 1)) / (eigenvec[[1]][[1]] *
            (1 - (1 - μ) * Exp[λ1 * T])) /. μ → mu /. ParmsTcA, T, Reals][[1]]
 ]
]
critTvec
pTcA = ListLinePlot[Transpose@{muvec, critTvec},
  PlotStyle → {RGBColor["#F564E3"]}, PlotLegends → {"T. colubriformis"}]
```

Out[•]=
```
{1.32793, 2.04413, 2.79791, 3.59145, 4.42699, 5.30688, 6.23353, 7.20947, 8.23735,
 9.32001, 10.4605, 11.6621, 12.9287, 14.2645, 15.6747, 17.1653, 18.7435, 20.4184}
```

Out[•]=
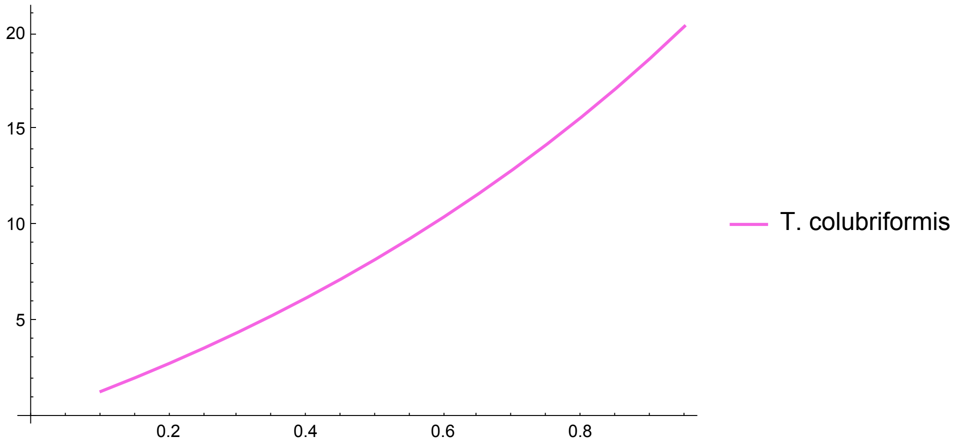


OcircNZ

In[•]:=
```
ParmsOc = {β → 1080.4 / 365, Nhost → 0.0015,
   μh → 0.1 / 365, μa → 40.9 / 365, λ → 4380 / 365, μl → 8.49 / 365};
critTvec = {};
muvec = {};
For[mu = 0.1, mu ≤ .99, mu += .05,
 AppendTo[muvec, mu];
 AppendTo[critTvec, T /. NSolve[(Exp[λ2 * T] - 1) / (1 - Exp[λ1 * T]) ==
        (eigenvec[[2]][[1]] * ((1 - μ) * Exp[λ2 * T] - 1)) / (eigenvec[[1]][[1]] *
           (1 - (1 - μ) * Exp[λ1 * T])) /. μ → mu /. ParmsOc, T, Reals][[1]]
 ]
]
critTvec
pOc = ListLinePlot[Transpose@{muvec, critTvec}, PlotStyle → RGBColor["#00BFC4"]]
```

Out[•]=
```
{0.123912, 0.19086, 0.261482, 0.336072, 0.414952,
 0.498478, 0.587042, 0.681079, 0.781068, 0.887543, 1.00109,
 1.12237, 1.2521, 1.39108, 1.54022, 1.7005, 1.873, 2.05893}
```

Out[•]=

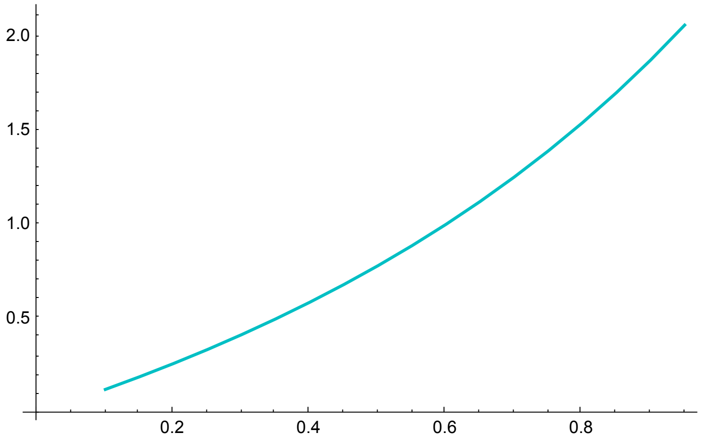


OcircUK

In[•]:=
```
ParmsOcUK = {β → 956.3 / 365, Nhost → 0.002,
   μh → 0.1 / 365, μa → 14.9 / 365, λ → 1752 / 365, μl → 11.12 / 365};
critTvec = {};
muvec = {};
For[mu = 0.1, mu ≤ .99, mu += .05,
 AppendTo[muvec, mu];
 AppendTo[critTvec, T /. NSolve[(Exp[λ2 * T] - 1) / (1 - Exp[λ1 * T]) ==
       (eigenvec[[2]][[1]] * ((1 - μ) * Exp[λ2 * T] - 1)) / (eigenvec[[1]][[1]] *
          (1 - (1 - μ) * Exp[λ1 * T])) /. μ → mu /. ParmsOcUK, T, Reals][[1]]
 ]
]
critTvec
pOcUK = ListLinePlot[Transpose@{muvec, critTvec},
  PlotStyle → {RGBColor["#00BFC4"], Dashed}, PlotLegends → {"O. circumcinta"}]
```

Out[•]=
```
{0.338299, 0.521076, 0.713886, 0.917529, 1.13288, 1.36093, 1.60272, 1.85947, 2.13247,
 2.42318, 2.73323, 3.0644, 3.41869, 3.79832, 4.20576, 4.64377, 5.11539, 5.62405}
```

Out[•]=

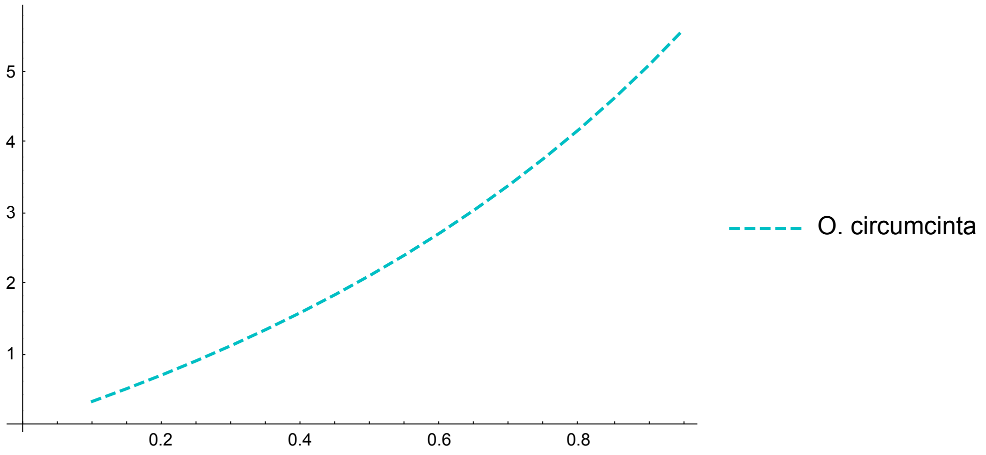


HcontNZ

In[•]:=
```
ParmsHc = {β → 584 / 365, Nhost → 0.0015,
   μh → 0.1 / 365, μa → 3.65 / 365, λ → 11 558.5 / 365, μl → 620.87 / 365};
critTvec = {};
muvec = {};
For[mu = 0.1, mu ≤ .99, mu += .05,
 AppendTo[muvec, mu];
 AppendTo[critTvec, T /. NSolve[(Exp[λ2 * T] - 1) / (1 - Exp[λ1 * T]) ==
        (eigenvec[[2]][[1]] * ((1 - μ) * Exp[λ2 * T] - 1)) / (eigenvec[[1]][[1]] *
           (1 - (1 - μ) * Exp[λ1 * T])) /. μ → mu /. ParmsHc, T, Reals][[1]]
 ]
]
critTvec
pHc = ListLinePlot[Transpose@{muvec, critTvec}, PlotStyle → RGBColor["#B79F00"]]
```

Out[•]=
```
{8.74957, 13.4909, 18.5152, 23.8587, 29.5644, 35.6853, 42.2863, 49.4494, 57.2793,
 65.9128, 75.5342, 86.3989, 98.8768, 113.532, 131.287, 153.818, 184.676, 233.906}
```

Out[•]=

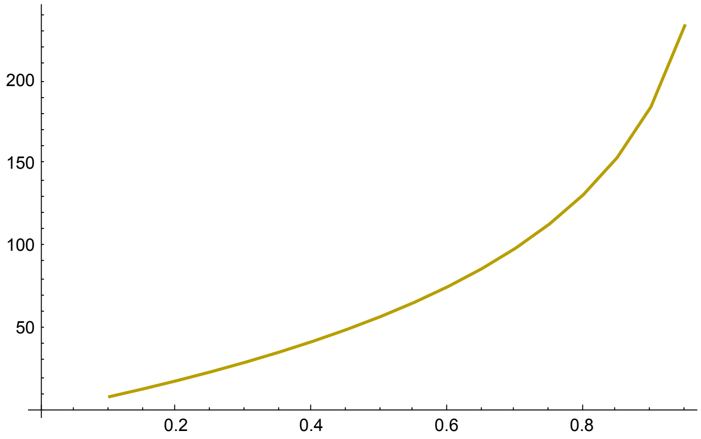


HcontAUS

In[•]:=
```
ParmsHcA = {β → 890.6 / 365, Nhost → 0.001,
   μh → 0.1 / 365, μa → 3.65 / 365, λ → 5501.8 / 365, μl → 230.36 / 365};
critTvec = {};
muvec = {};
For[mu = 0.1, mu ≤ .99, mu += .05,
 AppendTo[muvec, mu];
 AppendTo[critTvec, T /. NSolve[(Exp[λ2 * T] - 1) / (1 - Exp[λ1 * T]) ==
        (eigenvec[[2]][[1]] * ((1 - μ) * Exp[λ2 * T] - 1)) / (eigenvec[[1]][[1]] *
           (1 - (1 - μ) * Exp[λ1 * T])) /. μ → mu /. ParmsHcA, T, Reals][[1]]
 ]
]
critTvec
pHcA = ListLinePlot[Transpose@{muvec, critTvec},
  PlotStyle → {RGBColor["#B79F00"], Dotted}, PlotLegends → {"H. contortus"}]
```

Out[•]= {5.61202, 8.64613, 11.8548, 15.2594, 18.8858, 22.7647, 26.9342, 31.4411, 36.345, 41.7227, 47.6756, 54.3416, 61.9151, 70.6828, 81.0931, 93.9067, 110.577, 134.476}

Out[•]=

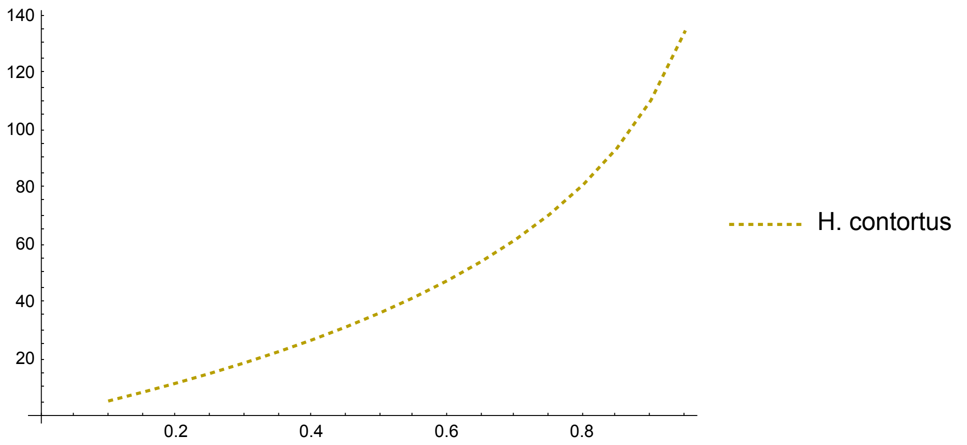


In[•]:=
```
Show[pTc, pTcA, pOc, pOcUK, pHc, pHcA, PlotRange → {0, 250},
 Frame -> True, FrameStyle → Directive[Black, 14]]
```

Out[•]=

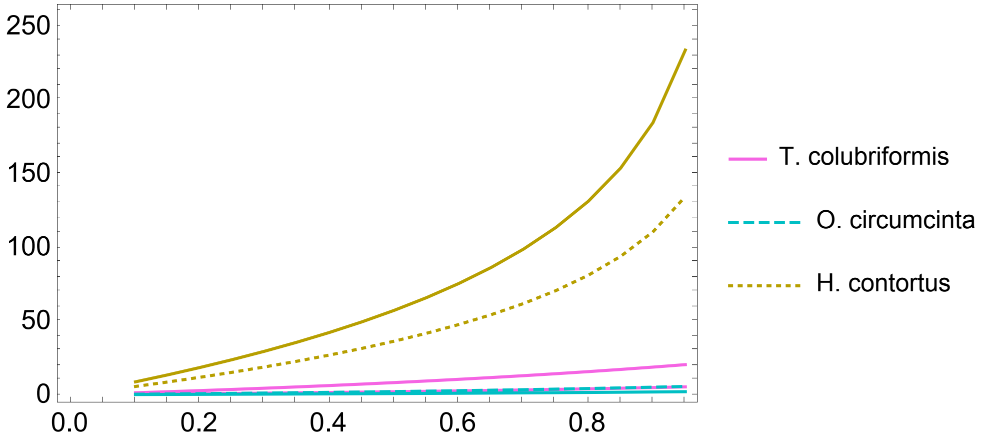

In[∘]:=
```
Show[pTcA, pOcUK, pHcA, PlotRange → {0, 150},
 Frame -> True, FrameStyle → Directive[Black, 14]]
```

Out[∘]=

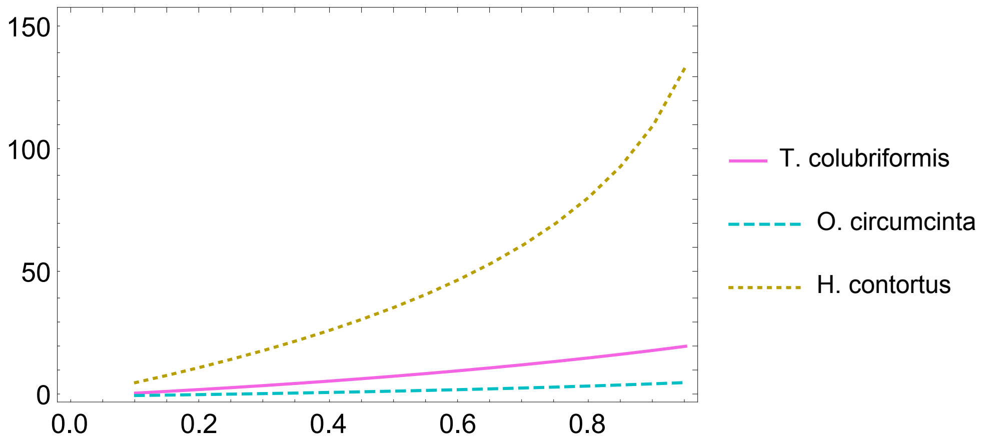


#### Categorizing Qualitative Behaviors

Reads in a csv file in which each row is a parameter set and determines the corresponding qualitative behavior: (1) decays even after evolution of resistance, (2) evolutionary rescue, (3) growth even before evolution of resistance, and (4) decay even without treatment (see Figure 3 in main text.)

```
LHS = Import["50grid.csv"];
critTvecbb = {};
critTvecaa = {};
determ = {};
dynvec = {};
For[idx = 2, idx ≤ Dimensions[LHS][[1]], idx += 1,
 parms = LHS[[idx]];
 Parmstemp = {β → parms[[7]], Nhost → parms[[6]], μh → parms[[9]], μa → parms[[4]],
   λ → parms[[5]], μl → parms[[8]], cov → 1, bb → 0.9, ab → 0.5, aa → 0.1};
 AppendTo[critTvecbb, T /. NSolve[(Exp[λ2 * T] - 1) / (1 - Exp[λ1 * T]) ==
          (eigenvec[[2]][[1]] * ((1 - μ) * Exp[λ2 * T] - 1)) / (eigenvec[[1]][[1]] *
             (1 - (1 - μ) * Exp[λ1 * T])) /. μ → cov * bb /. Parmstemp, T, Reals][[1]]
   ] ×
   AppendTo[critTvecaa, T /. NSolve[(Exp[λ2 * T] - 1) / (1 - Exp[λ1 * T]) ==
          (eigenvec[[2]][[1]] * ((1 - μ) * Exp[λ2 * T] - 1)) / (eigenvec[[1]][[1]] *
             (1 - (1 - μ) * Exp[λ1 * T])) /. μ → cov * aa /. Parmstemp, T, Reals][[1]]
   ] ×
   freq = parms[[3]];
 categ4 = If[critTvecbb[[idx - 1]] < 0, 4, 0];
 categ3 =
  If[critTvecaa[[idx - 1]] > 365 / freq && critTvecbb[[idx - 1]] > 365 / freq, 3, 0];
 categ2 = If[critTvecaa[[idx - 1]] < 365 / freq &&
    critTvecbb[[idx - 1]] > 365 / freq, 2, 0];
 categ1 = If[critTvecaa[[idx - 1]] < 365 / freq &&
    critTvecbb[[idx - 1]] < 365 / freq, 1, 0];
 AppendTo[dynvec, categ3 + categ2 + categ1 + categ4]
]
Export["50grid_results.csv", dynvec];
```
