## Supplementary figures and images for "Eco-evolutionary dynamics of anthelmintic resistance in soil-transmitted helminths"

### SI5 Additional Figures 1

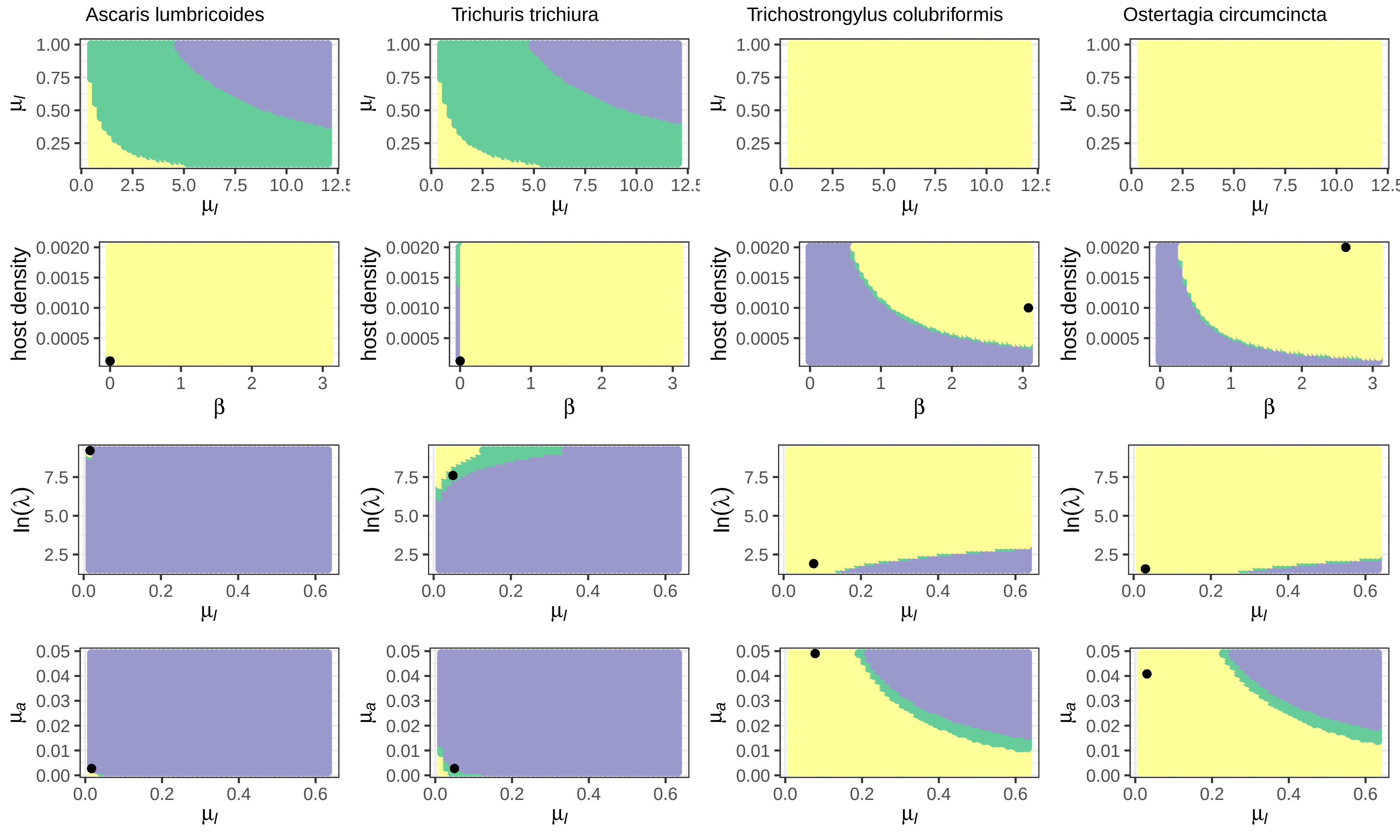

### SI6 Additional Figures 1

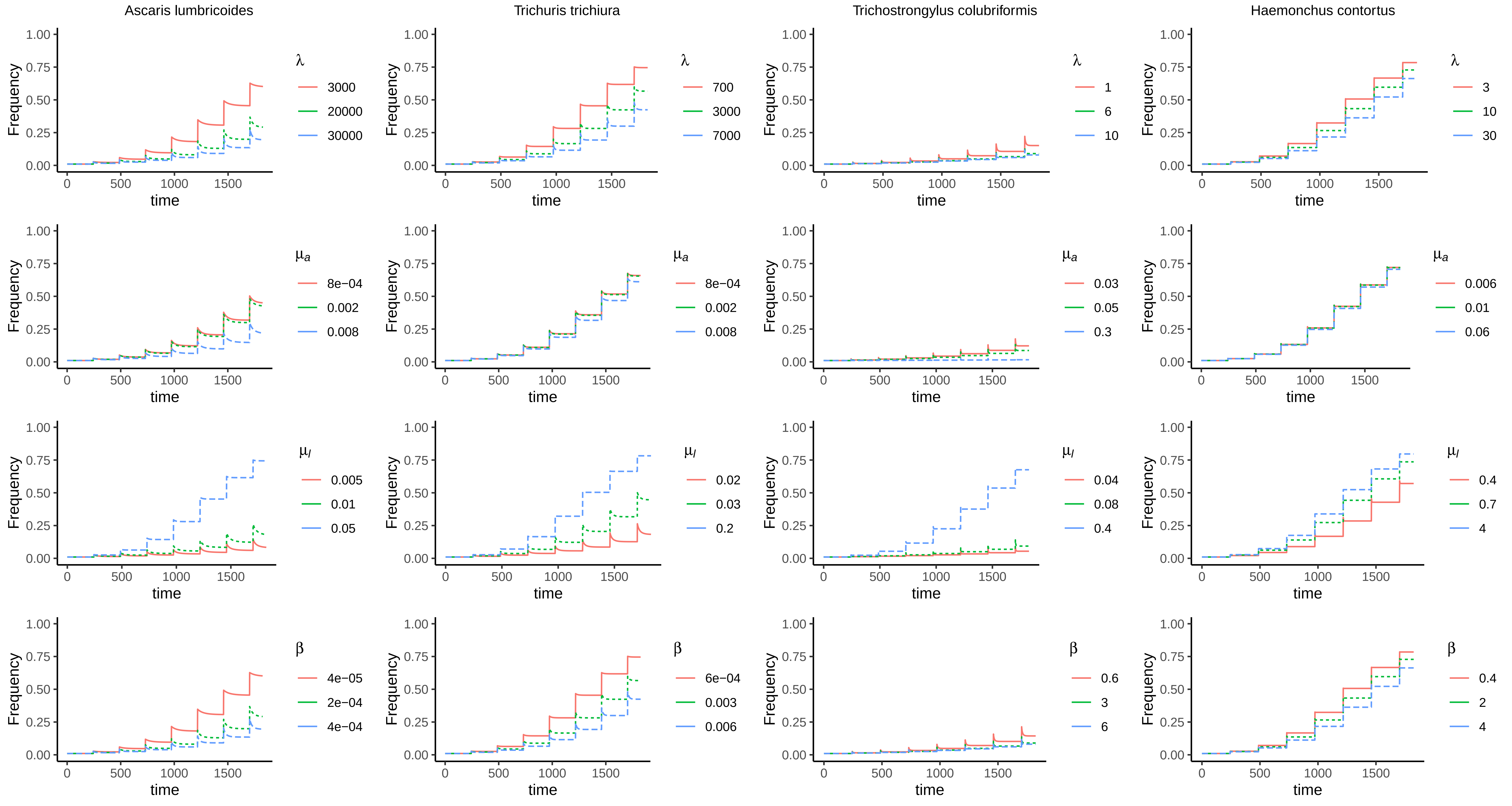
